# Sex-biased proteomic response to tomato spotted wilt virus infection of the salivary glands of *Frankliniella occidentalis,* the western flower thrips

**DOI:** 10.1101/2022.07.18.500439

**Authors:** Swapna Priya Rajarapu, Sulley Ben-Mahmoud, Joshua B. Benoit, Diane E. Ullman, Anna E. Whitfield, Dorith Rotenberg

**Affiliations:** Department of Entomology and Plant Pathology, North Carolina State University, Raleigh, North Carolina, 27695, USA; Department of Entomology and Nematology, University of California, Davis, Davis, California, 95616, USA; Department of Biological Sciences, University of Cincinnati, Cincinnati, Ohio, 45221, USA

**Author notes:** Corresponding author: Department of Entomology and Plant Pathology, North Carolina State University, Raleigh, North Carolina, 27695, USA.

**Keywords:** *Bunyavirales*, orthotospovirus, Thysanoptera, virus infection, saliva

## Abstract

Successful transmission of tomato spotted wilt virus (TSWV) by *Frankliniella occidentalis* requires robust infection of the salivary glands (SGs) and virus delivery to plants during salivation. Feeding behavior and transmission efficiency are sexually-dimorphic traits of this thrips vector species. Proteins secreted from male and female SG tissues, and the effect of TSWV infection on the thrips SG proteome are unknown. To begin to discern thrips factors that facilitate virus infection of SGs and transmission by *F. occidentalis*, we used gel- and label-free quantitative and qualitative proteomics to address two hypotheses: (i) TSWV infection modifies the composition and/or abundance of SG-expressed proteins in adults; and (ii) TSWV has a differential effect on the male and female SG proteome and secreted saliva. Our study revealed a sex-biased SG proteome for *F. occidentalis,* and TSWV infection modulated the SG proteome in a sex-dependent manner as evident by the number, differential abundance, identities and generalized roles of the proteins. Male SGs exhibited a larger proteomic response to the virus than female SGs. Intracellular processes modulated by TSWV in males indicated perturbation of SG cytoskeletal networks and cell-cell interactions (basement membrane, BM and extracellular matrix proteins, ECM), and subcellular processes consistent with a metabolic slow-down under infection. Several differentially-abundant proteins in infected male SGs play critical roles in viral life cycles of other host-virus pathosystems. In females, TSWV modulated processes consistent with tissue integrity and active translational and transcriptional regulation. A core set of proteins known for their roles in plant cell-wall degradation and protein metabolism were identified in saliva of both sexes, regardless of virus infection status. Saliva proteins secreted by TSWV-infected adults indicated energy generation, consumption and protein turnover, with an enrichment of cytoskeletal/BM/ECM proteins and tricarboxylic acid cycle proteins in male and female saliva, respectively. The nonstructural TSWV protein NSs - a multifunctional viral effector protein reported to target plant defenses against TSWV and thrips - was identified in female saliva. This study represents the first description of the SG proteome and secretome of a thysanopteran and provides many candidate proteins to further unravel the complex interplay between the virus, insect vector, and plant host.

## 1. INTRODUCTION

The order Thysanoptera is exclusively composed of thrips, a diverse group of tiny, fringe-winged insects. *Frankliniella occidentalis* (Suborder Terebrantia, Family Thripidae, Subfamily Thripinae), commonly known as the western flower thrips, is a destructive, polyphagous plant pest that causes damage directly by extensive feeding and oviposition on plant parts and indirectly by serving as the principal thrips vector of orthotospoviruses, some of the most economically-devastating viruses of food, fiber and ornamental crops (reviewed in Reitz et al., 2020). Due to its wide host range, worldwide distribution and ability to transmit eight of the 15 officially-described species of orthotospoviruses, *F. occidentalis* has been considered a ‘supervector’ that threatens global food security (reviewed in Gilbertson et al., 2015). Unlike other piercing-sucking insects, phytophagous thrips, for the most part, are mesophyll-feeders that consume entire cell contents, causing silvering of the leaf tissue (Kindt et al., 2003). The mouthparts of thrips are asymmetrical and adapted for ‘punch and suck’ feeding behaviors, whereby they form a mouth cone with two maxillae, one complete mandible and labium. The maxillary tubes fuse to form a sucking tube with a sub-terminal aperture (Hunter and Ullman, 1992). The *F. occidentalis* feeding behavior on plant tissue exhibits three electropenetrogram waveforms: non-ingestion probing, short-ingestion, and long-ingestion waveforms (Harrewijn et al., 1996; Stafford et al., 2011). Intermittent salivation is proposed to occur during non-ingestion (Stafford et al., 2011) and short-ingestion probes (Harrewijn et al., 1996; Stafford et al., 2011).

The salivary glands (SGs) of thrips consist of ovoidal principal and tubular secondary glands (Ullman et al., 1989). The principal glands are composed of eight large, binucleated, and loosely aggregated cells, whereas the tubular glands are composed of flat and mononucleated cells (Ullman et al., 1989). The tubular glands are connected to the anterior midgut and are proposed to deliver midgut digestive enzymes into the saliva (Ullman et al., 1989). The SG transcriptome of *F. occidentalis* comprises transcripts identified in other piercing sucking insects (Stafford et al., 2014). Protein-encoding sequences identified in *F. occidentalis* salivary gland transcriptome include proteases, carbohydrate degrading enzymes, calcium binding proteins, detoxification enzymes, and uncharacterized proteins (Stafford-Banks et al., 2014). Additionally, a comparative analysis of RNAseq datasets obtained from salivary gland tissue (Stafford-Banks et al., 2014)and whole-bodies of adult *F. occidentalis* revealed 123 transcripts enriched in expression in female and male SGs. These include transcripts encoding lipases, mannanases, deoxyribonucleases, and over half were uncharacterized in other metazoans, or putatively novel, *F. occidentalis* proteins (Rotenberg et al., 2020), several of which were predicted to be released into the plant host via saliva. Collectively, the proteins are hypothesized to play roles in host feeding and thrips-plant interactions.

Thrips SGs also play a defining role in the transmission of orthotospoviruses to plants. *Tomato spotted wilt orthotospovirus* is a member of the genus *Orthotospovirus* (Order *Bunyavirales*, Family *Tospoviridae*, TSWV), and is the most widely occurring orthotospovirus around the globe. It has served alongside *F. occidentalis* as the model system for molecular studies of thrips vector – orthotospovirus interactions (reviewed in Rotenberg and Whitfield, 2018). TSWV is a circulative-propagative virus that replicates in and traverses different thrips tissues (Montero-Astúa et al., 2016; Ullman et al., 1993; Whitfield et al., 2005) including the salivary glands. TSWV acquisition by larval thrips is required for thrips vector competency (Ullman et al., 1992a; Wijkamp and Peters, 1993) and adult thrips are transmission-competent only if virus is acquired as larvae. Adult thrips that ingest virus particles can sustain a midgut infection, but it does not lead to systemic infection (i.e., entry into the salivary glands) nor plant inoculation (Ullman et al., 1992b). Adults are the primary vehicles of TSWV inoculation and spread as they are winged and move plant-to-plant in the landscape.

The dissemination route of TSWV in *F. occidentalis* begins with attachment of virions to the anterior midgut epithelial cells, whereby the virus enters, replicates and disseminates cell-to-cell along a dedicated route from the midgut to the principal SGs, in parallel with the development of larvae to adults, where it first breaches the SGs during the late second instar larval stage (Montero-Astúa et al., 2016; Nagata et al., 1999; Ullman et al., 1993, 1992b). Virus is retained during the pupal stages, albeit at a lower abundance as compared to the larval stages (reviewed in Rotenberg and Whitfield, 2018). After adult eclosion, virus titers steadily increase in *F. occidentalis* bodies (Rotenberg and Whitfield, 2018), most notably due to TSWV replication in the SGs, a prerequisite to plant inoculation (Ullman et al., 1992b; Wijkamp et al., 1995, p. 1995). Virus accumulation is observed in both the primary and tubular glands (Badillo-Vargas et al., 2019; Montero-Astúa et al., 2016). Once infected, *F. occidentalis* adults can transmit the virus throughout their lifetime with a higher efficiency within 48 hours of their emergence (Rotenberg et al., 2009). Due, in part, to the difference in feeding behaviors, male *F. occidentalis* are more efficient vectors of TSWV (Rotenberg et al., 2009; Stafford et al., 2011; Van et al., 1998; Wetering et al., 1999). On plant tissue, feeding by females causes the most damage (necrotic scars) relative to feeding by males (water-soaked lesions) (Wetering et al., 1998; Stafford et al., 2011). Males typically engage in non-feeding behaviors and fewer probes of short duration, whereas females exhibit more longer-lasting, non-ingestion and short-ingestion probing behaviors compared to males (Stafford et al., 2011). Upon TSWV infection, the difference between males and females widens, wherein only infected males probe significantly more than their non-infected counterparts (Stafford et al., 2011).

Although inoculation efficiency, feeding behaviors, and tissue damage differ significantly between the sexes of *F. occidentalis*, there is a dearth of knowledge about proteins secreted from male and female SG tissues, and the effect of TSWV infection on the thrips SG proteome is unknown. Our study addressed two hypotheses: (a) TSWV infection modifies the composition and/or abundance of SG-expressed proteins in *F. occidentalis* adults; and (b) TSWV has a differential effect on the male and female SG proteome and secretome (saliva protein identification). We addressed these hypotheses using gel and label free quantitative proteomic experiments to 1) catalog the repertoire of SG-expressed proteins in male and female *F. occidentalis;* 2) determine the effect of TSWV on the diversity and abundance of SG-expressed proteins in male and females; and to 3) profile the proteins secreted in the saliva of male and female thrips exposed and non-exposed to TSWV. Our study provides proteomic evidence of sex-biased tissue expression and secretion (saliva) of *F. occidentalis* SGs, and revealed a sex-specific response to TSWV infection. To the best of our knowledge, this study represents the first direct protein-based assessment of the SG proteome and secretome of a thysanopteran.

## 2. MATERIAL AND METHODS

### 2.1. Insects, virus and plants

A colony of *Frankliniella occidentalis* (Pergande), originally started from an isolate from Kamilo Iki Valley, Oahu, Hawaii, was maintained on *Phaseolus vulgaris* green bean pods in deli cups covered with a ventilated lid as described previously (Ullman et al., 1992b) at 24^°^C with a daylight cycle of 14L:10D. The TSWV isolate MT2 was maintained on *Emilia sonchifolia* by thrips-mediated inoculations in the greenhouse (Badillo-Vargas et al., 2012). Infected leaf tissue from symptomatic plants was used to prepare inoculum for mechanical, rub inoculation (Ullman et al., 1993) of two-week-old plants of *Datura stramonium,* which were maintained in a growth chamber at 26^°^C until virus symptoms appeared. These plants served as the source of inoculum for experiments, and with each biological replicate, new batches of inocula were generated. Mock inoculations with inoculation buffer (10mM sodium sulfite in distilled water) were carried out similarly and served as the healthy controls. The optimal day for ≥ 90% acquisition efficiency by the first instar larvae was identified as 11-12 days post rub-inoculation.

### 2.2. Virus acquisition and salivary gland dissections

The design of the SG proteome experiment consisted of two factors: sex [males, M and females, F] and virus treatment [TSWV-exposure (V) and no virus exposure (NV)]. The experiment was repeated to generate four biological replicates (independent repetitions), i.e., new virus inocula, thrips cohorts, dissection events, and other materials. Due to the minute size of thrips, and likewise SGs, to obtain enough protein, each biological replicate was generated from a pool of SGs collected from three to eight trials of the virus acquisition assay. For TSWV acquisitions, larvae emerging within 17 hours from female-impregnated green bean pods were given a 24-hour acquisition access period (AAP) on young detached leaves from mock or TSWV-infected *Datura stramonium* leaves in leaf bouquets as previously described (Badillo Vargas et al., 2012) and contained in deli cups securely sealed with vented lids fitted with nylon mesh (100 micron, Elko Filtering Co.). The AAPs were carried out at 30^°^C in an incubator. Infections rates were determined on subsamples of ten 72-hour-old larvae (L2 stage) using *F. occidentalis* actin (reference gene) and TSWV-N gene-specific primers by quantitative real-time reverse transcription PCR or by PCR as described previously (Schneweis et al., 2017). Larvae fed on mock leaves were also tested and confirmed to be TSWV-negative. Larval cohorts with ≥ 70% infection rates were reared to adulthood for SG and saliva collection. Mock and infected larval cohorts were reared to adulthood on green bean pods.

Preliminary experimentation determined that for males and females, respectively, 200 and 100 paired SGs was the optimal number for a single gel-free nano liquid chromatography tandem mass spectrometry run yielding ∼2000 proteins. Newly emerged adults (24 - 48 hours after eclosion) were dissected in 1X phosphate buffer saline (PBS) with in-house dissection tools. Separate dissection tools were used for each treatment to prevent cross contamination, and surfaces were cleaned between treatments. Due to the duration of time required to dissect the necessary numbers of SGs, dissections were performed in alternating batches of NV and V males and females to minimize any possible effect of lapsed time on downstream statistical analyses. Dissected SGs were collected by a capillary needle along with 1X PBS buffer and centrifuged at 0.4 x g for two minutes. The supernatant was discarded and tissues were stored at −80^°^C and sent to the Duke Center for Genomics and Computational Biology - Proteomics and Metabolomics Core Facility for protein isolation, sample preparation, and quantitative proteomics through data processing and normalization. Refer to **Supplementary Fig. S1** for a flow chart of primary approaches and downstream analyses and data visualizations.

### 2.3. Protein isolation and sample preparation

The SG samples were treated with 200 µL of lysis buffer (8 M urea in 50 mM ammonium bicarbonate) and tissues were sonicated with an ultrasonic homogenizer with alternative washes to avoid contamination. Tissue lysate was centrifuged at 10000 x g for 10 minutes at 4^°^C. Total protein in an average 11 µL supernatant was estimated by the Pierce BCA assay following the manufacturer’s protocol for microplate assay (Thermofisher). An average of 6.1 µg (S.D. ± 0.7) of total protein was obtained from all the samples. Samples were supplemented with SDS for a final concentration of 3% for digestion and spiked with undigested casein at a total of either two or four pmol. Samples were then reduced with 10 mM dithiothreitol for 30 minutes at 80^°^C and alkylated with 25 mM iodoacetamide for 30 minutes at room temperature. Next, they were supplemented with a final concentration of 1.2% phosphoric acid and 2061 μL of S-Trap (Protifi) binding buffer (90% Methanol/100mM Tetraethylammonium bromide). Proteins were trapped on the S-Trap, digested using 32 ng/μL sequencing grade trypsin (Promega) for one hour at 47^°^C, and eluted using 50 mM Triethyl ammonium bicarbonate (TEAB), followed by 0.2% FA, and lastly using 50% acetonitrile with 0.2% formic acid. All samples were then lyophilized to dryness and resuspended in 40 μL 1% trifluoroacetic acid in 2% acetonitrile containing 12.5 fmol/μL yeast alcohol dehydrogenase.

### 2.4. Quantitative analysis of salivary gland tissues using nano liquid chromatography tandem mass spectrometry (LC-MS/MS)

Quantitative LC-MS/MS was performed on two μL (5%) of each sample using a nanoAcquity Ultra Performance Liquid Chromatography system (Waters Corp) coupled to a Thermo Orbitrap Fusion Lumos high resolution accurate mass tandem mass spectrometer (Thermo) via a nanoelectrospray ionization source. Briefly, the sample was first trapped on a Symmetry C18 20 mm × 180 μm trapping column (five μL/min at 99.9/0.1 v/v water/acetonitrile), after which the analytical separation was performed using a 1.8 μm Acquity HSS T3 C18 75 μm × 250 mm column (Waters Corp.) with a 90-min linear gradient of 5 to 30% acetonitrile with 0.1% formic acid at a flow rate of 400 nanoliters/minute with a column temperature of 55^°^C. Data collection on the Fusion Lumos mass spectrometer was performed in a data-dependent acquisition (DDA) mode of acquisition with a r=120,000 (@ m/z 200) full MS scan from m/z 375 – 1500 with a target Automatic Gain Control (AGC) value of 4e5 ions. MS/MS scans were acquired at Rapid scan rate (Ion Trap) with an AGC target of 1e4 ions and a max injection time of 100 milliseconds. The total cycle time for MS and MS/MS scans was two seconds. A 20s dynamic exclusion was employed to increase depth of coverage. The total analysis cycle time for each sample injection was approximately two hours. A statistical process quality control (SPQC) pool was created by taking five microliter from each sample, which was run periodically throughout the acquisition period.

### 2.5. Differentially abundant proteins in salivary glands

Data from UPLC-MS/MS were imported into Proteome Discoverer 2.4 (Thermo Scientific Inc.) and were aligned based on the accurate mass and retention time of detected ions (“features”) using the Minora Feature Detector algorithm in Proteome Discoverer. Relative peptide abundance was calculated based on area-under-the-curve (AUC) of the selected ion chromatograms of the aligned features across all runs. The MS/MS data were searched against the *F. occidentalis* (Focc_OGS.pep.v1, Rotenberg et al., 2019) and TSWV protein sequences (AAF80981.1 (G_N_/G_C_ glycoprotein precursor); AAL55403.1 (RNA-dependent RNA polymerase); ANG55707.1 (precursor G_C_, partial); and in house clones for G_N_, NSs, NSm, and N) and an equal number of reversed-sequence “decoys” for false discovery rate (FDR) determination. Mascot Distiller and Mascot Server (v 2.5, Matrix Sciences) were used to produce fragment ion spectra and to perform database searches. Database search parameters included fixed modification on Cys (carbamidomethyl) and variable modifications on Meth (oxidation), Asn and Gln (deamidation).

Peptide Validator and Protein FDR Validator nodes in Proteome Discoverer were used to annotate the data at a maximum 1% protein false discovery rate. Missing values were imputed after sample loading and trimmed-mean normalization in the following manner. If less than half of the values were missing in a treatment group, values were imputed with an intensity derived from a normal distribution defined by measured values within the same intensity range (20 bins). If greater than half of the values were missing for a peptide in a group and a peptide intensity is greater than 5e6, then it was concluded that peptide was misaligned and its measured intensity was set to 0. All remaining missing values were imputed with the lowest 5% of all detected values. Data were then subjected to a robust trim mean normalization excluding the highest and lowest 10% of the measured signals. Data were then rolled up from the peptide level to the protein level by summing all of the peptide intensities belonging to the specific protein. In order to assess technical reproducibility, we calculated the coefficient of variation (%CV) in the normalized abundance values for each protein across three injections of an SPQC pool that were interspersed throughout the run. To assess biological plus technical variability, %CVs were measured for each protein across the individual replicates of each treatment group. Protein intensities across all the samples were normalized to female SG protein abundances as male SG protein had lower signal than female SG proteins. The dataset of normalized protein abundance values was received from the proteomic facility, and then we identified differentially abundant proteins by the R package MSstats (v.3.4) (Choi et al., 2014). A linear model including sex and virus treatment main effects, i.e., non-exposed males (M-NV), TSWV-infected males (M-V), non-exposed females (F-NV) and TSWV-infected females (F-V), was fitted for every protein by MSstats and the interaction term of sex*treatment was examined by analysis of orthogonal contrasts. P-values were calculated for each pairwise comparison (eg. M-NV vs F-NV) post hoc, and adjusted p-values were calculated using the Benjamini and Hochberg method (Benjamini and Hochberg, 1995) to control the false discovery rate (FDR). Proteins with an FDR or P_adj_ < 0.05, and with more than one unique peptide that identified the protein, were considered differentially abundant proteins (DAPs).

Proteins were annotated against a non-redundant database (E-value cut-off = 10^-5^) in OmicsBox v2.1.14 (Götz et al., 2008). Gene ontology (GO) terms were assigned to all the proteins identified in the proteome with an E-value hit filter of 10^-3^ in OmicsBox. Enriched GOs within the DAPs between male and female SGs were identified by a two-tailed Fisher’s exact test with an FDR cut-off of 0.05 in OmicsBox (Al-Shahrour et al., 2004). In addition, a STRING analysis (STRING v11.5, https://string-db.org/, (Szklarczyk et al., 2017) was performed on DAPs between V and NV treatments to predict physical and functional protein-protein interaction (PPI) networks based on the provided *Drosophila melanogaster* proteome.

The TSWV peptides identified in the proteome were queried against the FOCC reference in NCBI (Blastp) with default parameters for short sequences, and thrips virome databases (PRJNA637687: Chiapello et al., 2021; PRJNA614937: Thekke-Veetil et al., 2020) using Blastp in local Blast+ with an E-value cut-off of 0.05 to identify any possible 100% identical matches to any other thrips viruses.

### 2.6. Prediction of secretome *in silico*

To identify SG proteins putatively secreted in the saliva, *in silico* analysis of the entire SG proteome was performed to predict up to three protein features: presence of a signal peptide, absence of transmembrane domains, and probable extracellular localization of the protein. Signal peptides were predicted by signalP (v.5.0) (Teufel et al., 2022) software against a eukaryotic database. Transmembrane helices were predicted with TMHMM (v.2.0) (Krogh et al., 2001; Sonnhammer et al., 1998). Proteins with more than 18 expected amino acids (ExpAA) within the predicted region were considered transmembrane. Cellular localization was predicted by DeepLoc (v.1.0) (Thumuluri et al., 2022) which uses the sequence information to predict its localization rather than homology-dependent prediction. As a stringent measure, proteins predicted to contain all three features were considered to be secreted. However, since there is also evidence of non-canonical proteins in saliva of hemipterans (reviewed in Sharma et al., 2014, those *F. occidentalis* SG proteins with only extracellular localization patterns were also considered putatively secreted into the saliva.

### 2.7. Collection of saliva from *F. occidentalis* adults

Since large numbers of thrips and feeding chambers were needed to obtain enough protein for discovery proteomics, 12 independent acquisition assays with >2000 insects per treatment (+/- TSWV) were conducted over the course of five months, which provide one pooled saliva sample per sex-treatment combination. Secreted saliva was collected from 24-48 hour old adults. Approximately, 3200 females and 2000 males per treatment were fed a sterile 3% sucrose diet with 0.4% resorcinol to aid in salivation (Chaudhary et al., 2015), sandwiched between parafilm layers over one fluid-ounce medicine cup to form a ‘feeding arena’. Diet volumes were calibrated on a per insect basis to achieve 1 µL of diet per insect placed in the arena, e.g. 100 µL for 100 adults in an arena. All cups were inverted to increase the probability of feeding by males. Feeding arenas with either males or females (NV and V) were maintained on the diet at 25^°^C with a daylight cycle of 14L: 10D for 24 hours. Diet which contained the secreted saliva was collected with a 200 µL sterile, filter-barrier tip in the laminar hood. Diet collected from the females was centrifuged at 400 x g for two minutes to sediment any eggs, and supernatant was stored at −80^°^C. Diet collected from the males was stored directly at −80^°^C. In total, 2.1 mL, 1.7 mL, 1.1 mL, and 1.2 mL of diet was collected from NV and V females and males, respectively. Based on the Bradford estimation of total protein of healthy adult saliva, females secreted an average of 14 μg/mL and males secreted an estimate of 2 μg/mL protein. This difference in the total protein secreted by males and females could be due to the difference in their feeding behaviors. Samples were freeze-dried to concentrate in the VirTis Model 24DX48 GPFD 35L EL-85 (SP scientific) freeze dryer. The tubes were vented and dried over 24 hours with 5^°^C-temperature decrements every eight hours starting from −30^°^C to 10^°^C. Samples were stored at −80^°^C and sent to the Duke Center for Genomics and Computational Biology – Proteomics and Metabolomics Core Facility for sample preparation and qualitative proteomics.

### 2.8. Qualitative analysis of secreted saliva proteins using nano liquid chromatography tandem mass spectrometry (LC-MS/MS)

Freeze dried saliva samples from the 12 independent acquisitions were resuspended in 100 µl of 8 M urea, sonicated with an ultrasonic homogenizer, and pooled to achieve sufficient amounts of protein per treatment. Samples were prepared for LC-MS/MS analysis following S-Trap^TM^ microspin column digestion protocol (PROTIFI). Briefly, the samples were supplemented with 20% SDS to a final concentration of 3%, reduced with Tris(2-carboxyethyl) phosphine, alkylated with Methyl methanethiosulfonate, acidified with phosphoric acid, and loaded onto S-traps for trypsin digestion. Proteins were eluted from the column and lyophilized to dryness. Samples were resuspended in 12 μL 1% trifluoroacetic acid in 2% acetonitrile containing 12.5 fmol/μL yeast alcohol dehydrogenase and one third of the resuspended sample was loaded onto the column. Qualitative LC-MS/MS was performed on four μL (5%) of each sample using a nanoAcquity Ultra Performance Liquid Chromatography system (Waters Corp) coupled to a Thermo Orbitrap Fusion Lumos high resolution accurate mass tandem mass spectrometer (Thermo) via a nanoelectrospray ionization source and a FAIMS pro Duo interface to increase the number of proteins detected.

Proteins in the secreted saliva were identified from the peptide sequence databases using Mascot search engine (v.2.5.1). Peptide spectra were searched against the *F. occidentalis* genome derived proteome database (Focc_OGS.pep.v1) and TSWV specific proteins with carbamidomethyl as the fixed modification and oxidation as the variable modification. Trypsin was specified as the proteolytic enzyme with a minimum of two missed cleavages. The qualitative analysis of the proteins was performed in the Scaffold viewer (v.4.9.0). Confidence of the protein identification was determined by the number of exclusive unique peptides per protein. For a comparative analysis between the samples, “exclusive unique spectrum count” was used to identify the proteins that are similar and unique between the samples. Proteins identified in both the SG proteome and secreted into the diet were considered “secreted-saliva” proteins. This saliva protein dataset was compared to a published *F. occidentalis* SG transcriptome (Stafford-Banks et al., 2014) and SG-enriched transcript sequences (Rotenberg et al., 2020). Saliva-secreted proteins were annotated in OmicsBox v2.1.14 (Götz et al., 2008). Enriched GO terms were identified by a two-tailed Fisher’s exact test with an FDR cut-off of 0.05 (Al-Shahrour et al., 2004) and the distribution of saliva proteins into GO categories (multilevel terminal nodes) was generated for each sex-treatment group separately with an E-value cut-off of 10^-3^. In addition, a STRING analysis (STRING v11.5, https://string-db.org/, (Szklarczyk et al., 2016) was performed to predicted physical PPI networks from the saliva protein datasets, with particular attention to proteins uniquely found in saliva of infected males and females.

### 2.9. Comparative analysis of *F. occidentalis* secreted saliva proteins with other piercing-sucking insect saliva proteins

Saliva proteins of *F. occidentalis* were compared with published saliva protein datasets for plant-feeding hemipterans. Proteins from the secreted saliva of representative species from each family within Hemiptera and an outgroup, *Tetranychus urticae* (Arachnida) - a herbivorous arthropod with a similar feeding strategy as *F. occidentalis* females (piercing and removal of mesophyll cell contents) and wide plant host range - were retrieved from NCBI based on the accession numbers provided in each respective study. Species compared included the potato aphid (*Macrosiphum euphorbiae,* 38 proteins) (Chaudhary et al., 2015), Asian citrus psyllid (*Diaphorina citri*, 211 proteins) (Wu et al., 2021), brown planthopper (*Nilaparvata lugens*, 49 proteins) (Huang et al., 2018), whitefly (*Bemisia tabaci*, 166 proteins) (Huang et al., 2021), green peach aphid (*Myzus persicae*, 171 proteins) (Guo et al., 2020) and two spotted spider mite (*Tetranychus urticae,* 95 proteins) (Jonckheere et al., 2016), A Blastp search was performed with the *F. occidentalis* saliva proteins as the query database with an E-value cut-off of 10 E^-5^ in NCBI Blast+ (v. 2.12) locally. Similarity was determined by matches with the highest bit-scores and lowest E-values.

### 2.10. Data availability

The raw proteomic data was deposited in Mass Spectrometry Interactive Virtual Environment (MassIVE) (https://massive.ucsd.edu/ProteoSAFe/static/massive.jsp) under the accessions MSV000089639_reviewer for the salivary gland proteome and MSV000089640_reviewer for the saliva proteome. Raw peptide sequences, modifications identified, retention times, and ion scores per protein identified in the SG samples are listed in **Supplementary File 1: Table S12**, and raw peptide sequences and other metrics per protein identified in secreted-saliva samples are provided in the **Supplementary File 2**. TSWV protein (GN, NSs, NSm, and N) sequences derived from in-house clone sequences for the TSWV MT2 isolate used in this study are provided in **Supplementary File 3**.

## 3. RESULTS

### 3.1. Analytical and biological reproducibility in protein abundance achieved for the *F. occidentalis* adult salivary gland peptide dataset

The dataset for SG tissues contained 27,041 peptide matches, and 1,486,468 MS/MS spectra were acquired for peptide sequencing. Following database searching and peptide scoring, annotation resulted in the quantification of 2,692 proteins identified by one to 364 unique peptides per protein. The mean %CV in the protein normalized abundance values for the SPQC pool samples was 15.3% for all proteins, an indication of technical reproducibility in the dataset. The average %CV in normalized abundance of proteins in the treatment samples, which captures both analytical and biological variability, was 27.6%, 24.8%, 22.3%, and 19.5% for the M-NV, M-V, F-NV, and F-V groups, respectively. A principal component analysis of all quantified proteins showed a tight clustering of the SPQC pool samples, and a clear differentiation between treatment groups (**Supplementary Fig. S2.A**), indicating biological reproducibility across the independent replications of the experiment. Overall, it appears that sex explained most of the variation (PC1, M vs F) in protein normalized abundance between treatment groups.

### 3.2. Metrics of the filtered salivary gland protein dataset

After removal of 28 contaminating and sample-processing proteins (egs., human keratin, yeast alcohol dehydrogenase) and *F. occidentalis* accessions identified with only one peptide (368), the final salivary gland dataset included 2,292 non-redundant proteins annotated as *F. occidentalis* proteins and three TSWV structural (N, G_N_/G_C_ glycoprotein precursor) and nonstructural (NSs) proteins in both sexes (**Supplementary File 1: Table S1**). The proteome included 6.8% uncharacterized proteins that are unique to *F. occidentalis*. The median number of unique peptides that identified a *F. occidentalis* protein was six (mean ∼ 11), with the majority of proteins (90%) identified by 2 to 24 unique peptides per protein. The median normalized abundance of protein across all identified thrips proteins (expressed as log10) was 5.382, with little variation in the median between treatment groups. However, the collection of proteins in male cohorts, regardless of virus infection status, tended to have five times higher minimum normalized abundance values (M-NV = 4.333; M-V = 4.318) compared to their female counterparts (F-NV = 3.656; F-V = 3.579), as well as a few more proteins in the top 1% of expressed proteins (**Supplementary Fig. S2.B**). The top ten most abundant *F. occidentalis* proteins in male and female salivary glands are listed in **Supplementary Fig. S2.C,** with lipophorin precursor as the most abundant protein across both the sexes, and vitellogenin among the most abundant proteins.

### 3.3. *In silico* analysis of salivary gland proteins reveals signatures of secretion

Canonical proteins predicted to be secreted into the saliva typically possess an N-terminal signal peptide, extracellular localization, and absence of transmembrane domain. Proteins with these features were independently identified in the SG proteome of *F. occidentalis*. We predicted 240 proteins (10.5%) with the presence of signal peptide, 2163 proteins without a transmembrane domain (94.3%), and 185 proteins with extracellular localization (8.2%). Among these, 123 (5.4%) proteins are predicted to contain all three features, providing strong evidence of secretion and possibly destined for saliva. Hemipteran proteins secreted in the saliva include non-canonical proteins (reviewed in Sharma et al., 2013), and as such, proteins containing predicted extracellular localization patterns (185) were considered candidates for secretion into the saliva (**Supplementary File 1: Table S2**).

### 3.4. Expression of *F. occidentalis* salivary gland proteins influenced by sex

Of the 2,292 salivary gland proteins in the final dataset, 51% and 52% were differentially-abundant between male and female thrips (*P* adj. ≤ 0.05, **Supplementary File 1: Table S3**) in the TSWV-exposed (V) and non-exposed (NV) treatment groups, respectively. Cross-examination of these sex-associated, differentially-abundant proteins (sDAPs) between treatment groups revealed an overlap of 930 proteins (by protein identifiers), while 245 and 268 proteins were uniquely associated with the V and NV treatments, respectively (**Supplementary Fig. S3**). Consistent between treatment groups, 51% of the sDAPs were up-regulated in salivary glands of females compared to males, and 48% were up-regulated in salivary glands of males compared to females.

### 3.5. Sex-specific enrichment of gene ontologies (GOs) associated with SGs

Depending on treatment groups, approximately 18.2% - 23.5% of the identified sDAPs were assigned GO terms (**Supplementary File 1: Table S4**). Irrespective of virus treatment, sDAPs in females had GO annotations that inferred protein metabolism, including ‘cellular nitrogen compound process’, ‘translation’, ‘ribosome’, and ‘RNA binding’, while sDAPs in males had GO annotations associated with enzymatic activities, such as ‘protein modification process’, and ‘mitochondria’. A GO enrichment analysis supported the finding that male sDAPs were underrepresented in processes and cellular components associated with protein translation (three categories, **Fig. 1A** and **B**) compared to the SG proteome at large, but were enriched in proteins associated with ‘lipid binding’ in the V treatment group (**Figure 1B**). Ten proteins within the ‘lipid binding’ GO were mostly represented by cell structural and endocytic pathway proteins including annexin-B9 & B10, sorting nexin 2 & lst; two spectrin-β chains; AP2-complex subunit; golgi phoshoprotein 3 homolog sauron; hydroxyphenyl pyruvate dioxygenase; and peroxisomal-acyl-coA oxidase. On the contrary, sDAPs in female SGs were enriched in several GO terms associated with protein translation regardless of virus treatment (**Fig. 1A** and **B**) compared to the total SG proteome, with some enriched GOs specific to V or NV treatment groups. Notably, sDAPs were enriched in the GO term ‘peptidase activity’ in the NV females only (*P* = 0.04), with approximately one-third of the proteins associated with the ubiquitin-mediated protein degradation pathway involving the 26S proteasome and ubiquitin carboxyl-terminal hydrolases. Four GO terms were enriched in V female SGs, including ‘hydrolase activity’ (*P* = 0.05), the category with the greatest number of proteins (59), a third of which are proteases and peptidases (catabolism), indication of increased protein turnover in female SGs.

**Figure 1.**
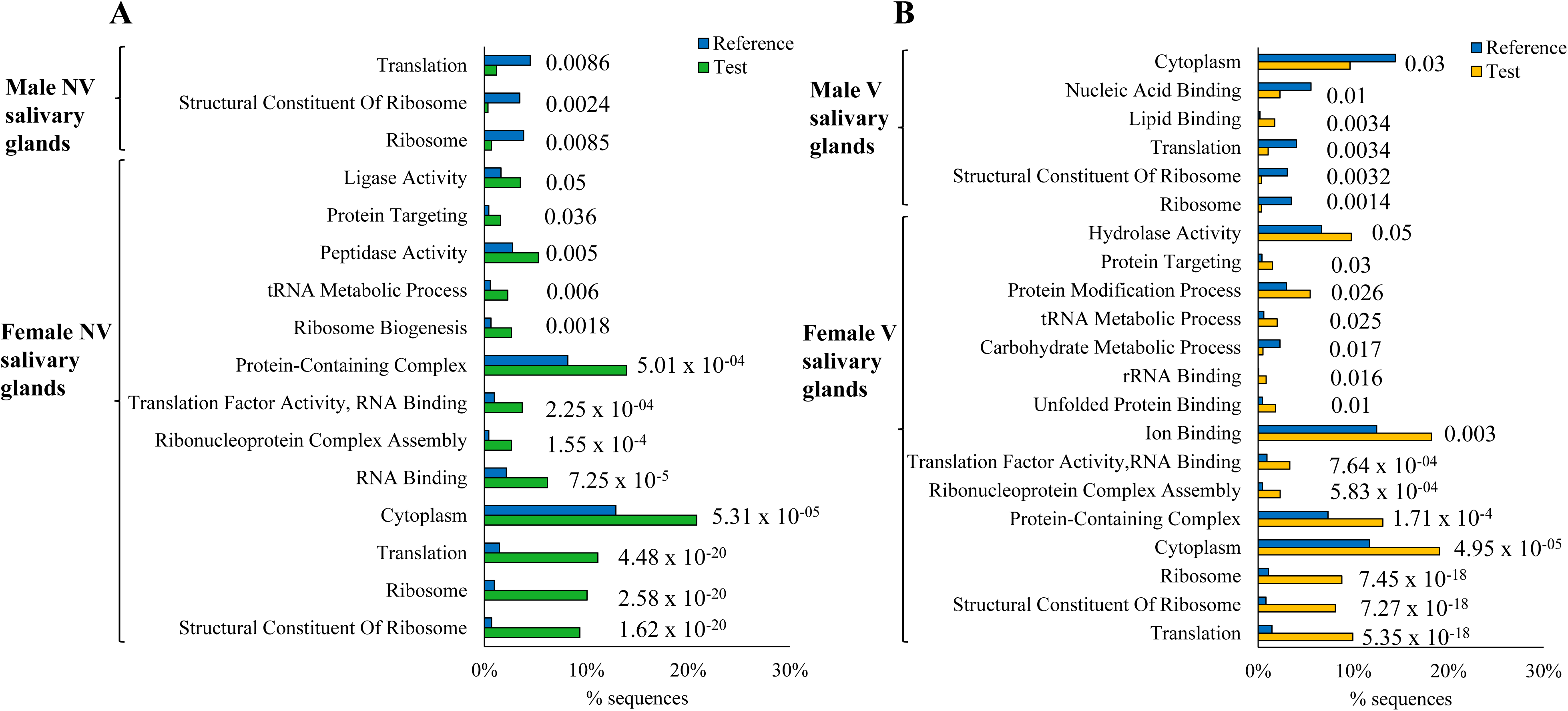
Enrichment of gene ontologies (GOs) associated with differentially abundant proteins (DAPs) in salivary glands (SGs) of male and female *Frankliniella occidentalis*. A two-tailed Fisher exact test (FDR cut-off of 0.05) was performed in OmicsBox v2.1.14 to compare sex-associated DAPs against the salivary gland proteome dataset (2,292 protein sequences). Comparisons were performed for each virus treatment cohort separately; non-exposed (NV) and tomato spotted wilt virus-infected (V). The FDR value of the enrichment is indicated above GO term bars (% of sequences enriched in each GO). “Reference” refers to the salivary gland proteome dataset with test set sequences removed. “Test” refers to enriched sex-associated DAPs for the respective sex-virus treatment combination. Refer to Supplementary File 1: Table S4 for the identities of GO-enriched SG proteins.

### 3.6. Sex-specific response of salivary glands to TSWV infection

The effect of TSWV infection on the adult SG proteome differed markedly between the sexes (**Fig. 2**). There were 2.6 times more virus-associated DAPs (vDAP) (58 vs 21) in male SGs compared to female SGs (**Fig. 2A and 2B**), and the proportion of down-regulated proteins within the sexes appeared to be greater in males compared to females (51.7% vs 31.8%). Overall, virus exposure had a larger effect on the proteome response of the male SGs compared to female SGs as determined by the absolute difference in normalized abundance of vDAPs between V and NV treatments (M = avg. (log10 value) = 5.253 (n = 58), F = avg.(log10 value) = 4.696 (n = 21), *P* = 0.0001) (**Fig. 2C**). In addition, the overlap in the vDAPs in female and male SGs was negligible (**Fig. 2D**) - only one down-regulated protein was shared between the sexes, putatively identified as a peroxiredoxin-4.

**Figure 2:**
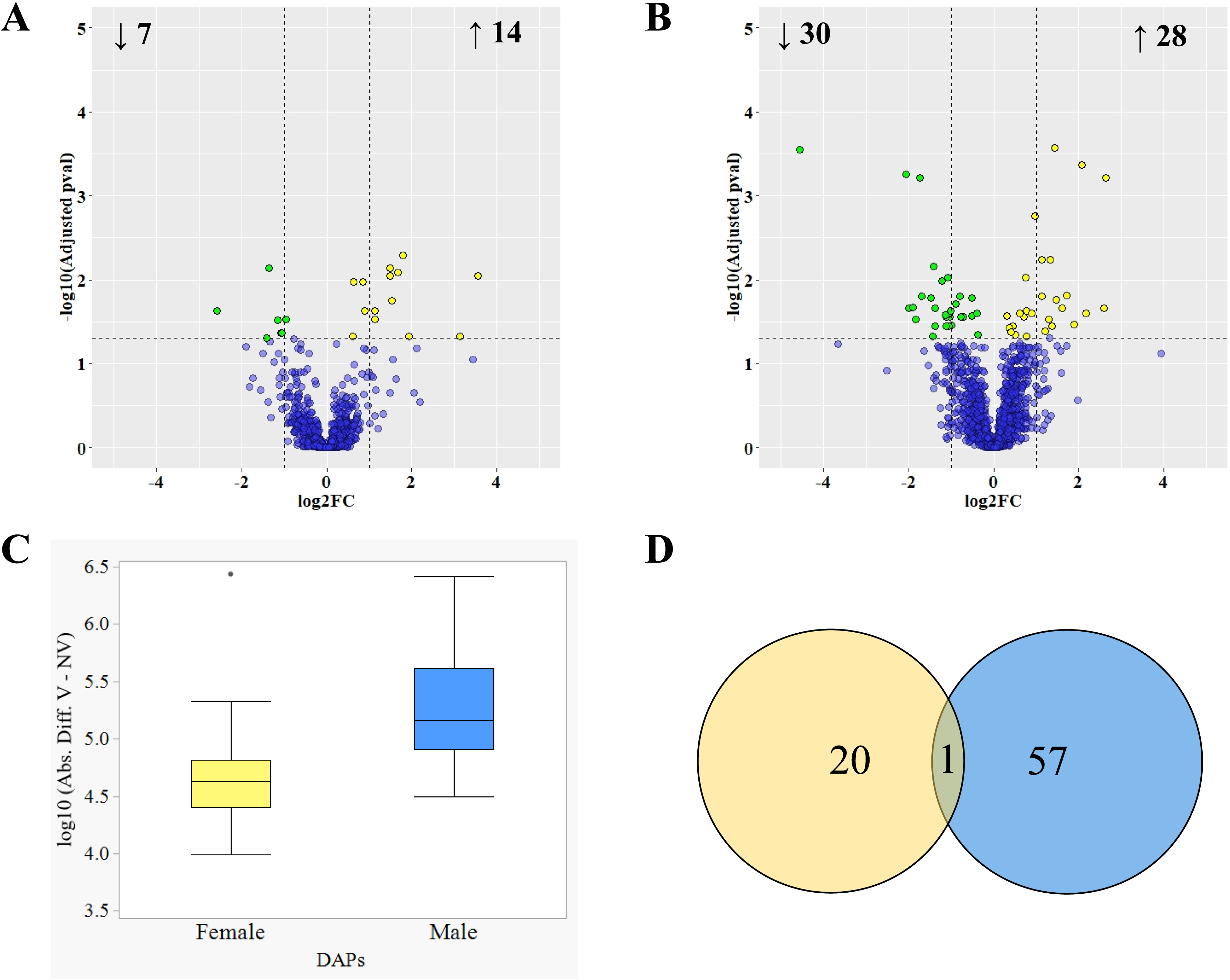
The differential effect of tomato spotted wilt virus (TSWV) infection on the salivary gland (SG) proteome of male and female *Frankliniella occidentalis*. (**A, B**) Volcano plots depicting magnitudes and direction of change in normalized abundance (log_2_ fold change) of proteins in SGs and associated levels of significance (adjusted p-value, log_10_) for pairwise comparisons between non-exposed (NV) and virus-infected (V) males (**A**) and females (**B**). The horizontal dashed line indicates the cut-off for statistical significance (0.05), and vertical dashed lines demarcate fold change values of ± 1.5. Yellow and green points indicate significantly up- and down-regulated proteins between virus treatments, i.e., TSWV-modulated proteins (*P*_adj_ < 0.05, each point represents n = four biological repetitions of the experiment). **(C)** Boxplots depicting the variation in the absolute difference in normalized protein abundance (log_10_-transformed) between virus treatments (V – NV) for proteins significantly modulated by virus infection. The mean magnitude of change (indicated as horizontal line in each boxplot) was significantly different between male and female SGs (P < 0.0001). **(D)** Venn diagram depicting the number of virus-modulated proteins unique and shared between the sexes.

The identities, fold change values (log2-transformed), and blastp annotations of the 78 vDAPs in SGs are presented in **Supplementary File 1**: **Table S5.** For the most part, the female-specific vDAPs with significant matches to known insect proteins inferred roles in tissue growth, neurogenesis, detoxification and/or hormone metabolism, carbohydrate- or actin-binding, proteolysis, translation (ribosomal subunits), RNA-binding and transcription regulation. Male- specific vDAPs inferred cytoskeletal, basement membrane and extracellular matrix proteins, diverse RNA-binding activities primarily associated with nuclear activities, mitochondrial activities, proteolysis, metal-ion binding critical for several catalytic activities, and intracellular transport of electrons, metal ions and proteins (e.g., sorting nexin).

Mapping the 78 vDAP sequences against a representative *D. melanogaster* proteome in STRING revealed 11 enriched molecular and subcellular activities perturbed by virus in male compared to female SGs (**Fig. 3**), and the proteins associated with these activities could be classified into four PPI networks. Enriched functions included ‘metal ion-binding’ (zinc, elemental sulfur, copper and manganese) - ‘transferase’ (N-acetyl transferases, including spermidine/spermine (SAT) transferase) and ‘hydrolase’ activities, ‘nucleotide-binding’ (for the most part, ‘RNA-binding’), transport (electrons, metal ions, and lysosomal) and ‘ATP-binding’. Many of these proteins play evolutionarily conserved activities in the ‘nucleus’ and ‘mitochondria.’ These virus-perturbed (UP or DOWN regulated) cellular activities in male SGs predicted known protein networks: i) cell-cell interactions (cytoskeleton, actin-binding, extracellular matrix (ECM), and basement membrane (BM) proteins)(most UP); ii) fatty acid synthesis (UP) and oxidation/electron transfer for respiration (DOWN); iii) proteolysis (UP) - proteasome (nucleus) and metallopeptidase - and electron carrier activity (mitochondrial) (DOWN); and iv) the largest protein network associated with RNA processing: small ribonucleoproteins (RNPs)(UP) occurring in the nucleus, specifically during the early and late spliceosome pathway (pre-mRNA splicing), ras GTPase-activating protein-binding protein (G3BP-type RNA-binding protein) associated with post-transcriptional modification of gene expression (UP), ATP-dependent RNA-helicases (DOWN), histone H3-H2 methylation (DOWN), biogenesis of ribosomal subunits (DOWN), as well as other mitochondrial activities (translation elongation factor and protein folding (both DOWN).

**Figure 3.**
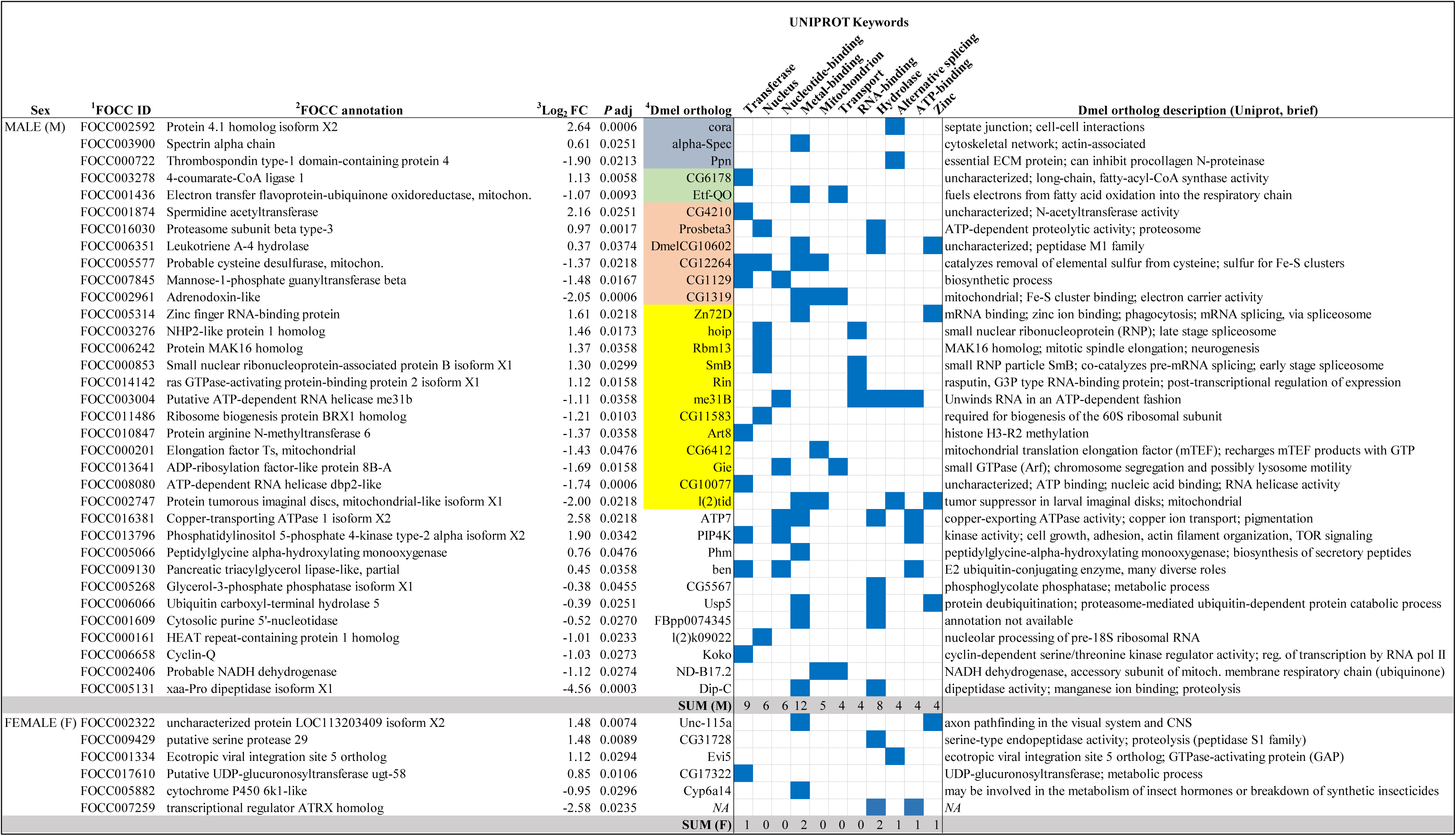
Sex-biased enrichment of predicted protein function and subcellular localization in salivary gland (SG) cells of tomato spotted wilt virus-infected adults of *Frankliniella occidentalis*. The dataset of TSWV-modulated proteins (78) were mapped to the *Drosophila melanogaster* proteome to identify putative orthologs and protein-protein interaction (PPI) networks using STRING analysis (https://string-db.org/, Szklarczyk et al., 2016). Depicted are the ^1^*F. occidentalis* proteins sequences (FOCC ID, Official Gene Set v1.1, Rotenberg et al., 2019) with associated NCBI or Uniprot ^2^accessions that were classified (blue squares) into 11 sex-enriched functional roles or compartments, i.e., keywords associated with the Uniprot annotations of the Dmel orthologs. ^3^Log_2_FC = log_2_-transformed fold change in the normalized abundance of the identified protein in SGs of virus-infected males and females compared to non-exposed counterparts, and associated adjusted p-values. ^4^Color-shading (gray, green, orange and yellow) indicates membership in one of four predicted physical-functional networks of proteins. Refer to Figure 4A for illustration and annotation of these four male-enriched PPI networks.

There was no apparent enrichment for particular processes in infected female SGs, which could be due, in part, to the nearly 3-fold fewer vDAPs in females compared to males. With regards to network association, central to the predicted “cell-cell interaction” network described above for male-enriched processes, was the vDAP in infected female SGs, integrin beta-PS (Dmel ortholog = mys). Integrin beta-PS is a transmembrane receptor protein for laminin, and it is associated with maintaining the integrity of the endodermis, and adhesion of midgut epithelia to surrounding visceral muscles (Brown et al., 2000). The predicted STRING PPI network of the SG vDAPs for both sexes combined is depicted in **Figure 4A**.

**Figure 4.**
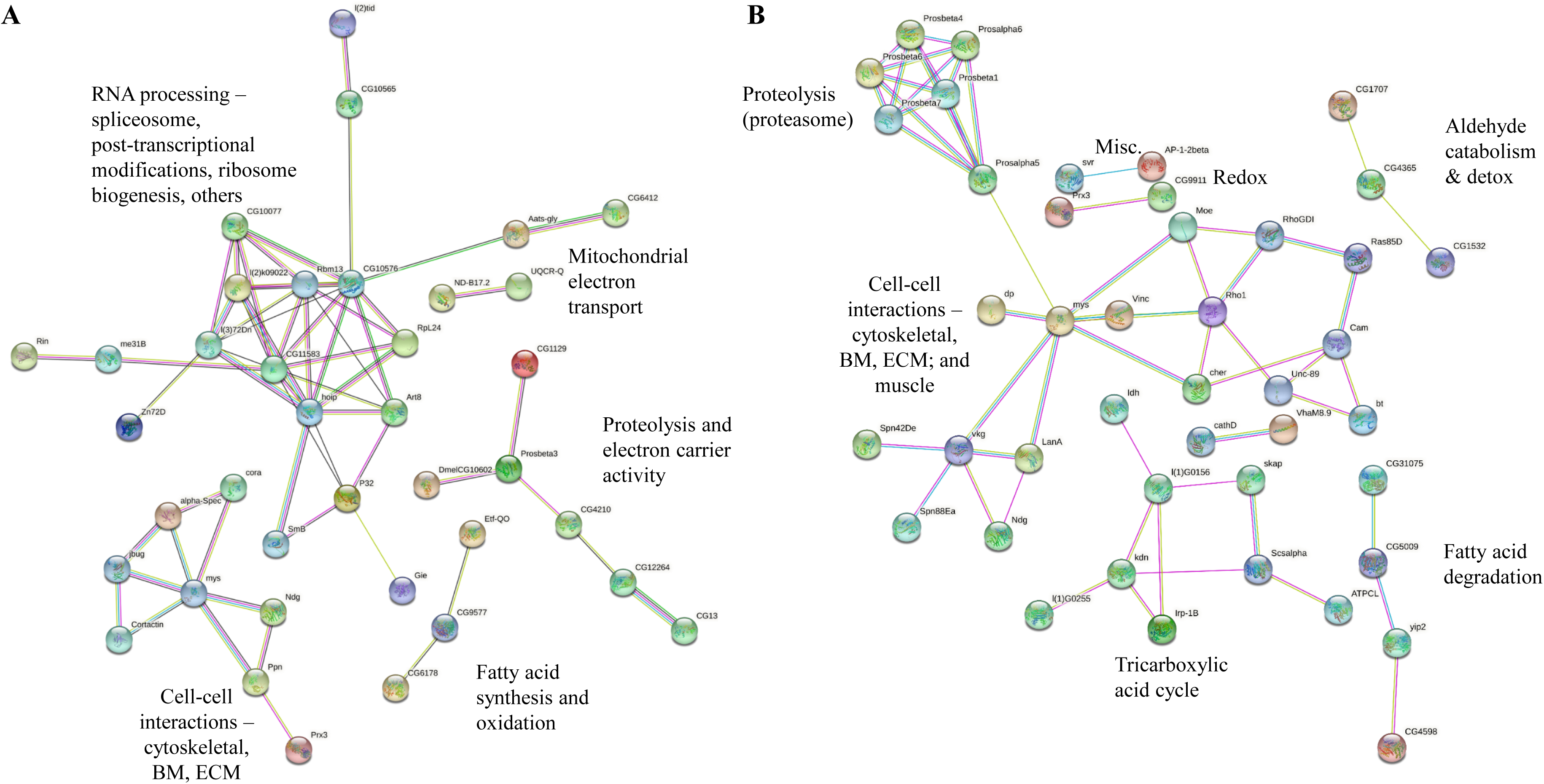
Predicted protein-protein interaction (PPI) networks associated with tomato spotted wilt virus (TSWV)-infected *Frankliniella occidentalis* adults. (**A**) Physical-functional PPI networks for differentially-abundant proteins in salivary glands (SG) in response to TSWV, and (**B**) physical PPI networks for SG-secreted saliva proteins unique to TSWV-infected adults. PPI networks were generated using STRING v11.5 (https://string-db.org/, Szklarczyk et al., 2016) on the software-provided *Drosophila melanogaster* (Dmel) proteome. Only proteins that mapped to the Dmel proteome and had a predicted interaction with at least one other protein are presented.

### 3.7. *Frankliniella occidentalis* saliva proteins were secreted and identified

Due to the minute size of thrips, and small amount of secreted saliva produced by thrips during feeding, our analysis was limited to qualitative comparisons (protein occurrence and raw spectral counts) between pooled samples for each sex-treatment combination. A total of 738 proteins from 15858 peptides and 82237 spectra were identified, and removal of contaminants, internal standards, and sample-processing proteins (trypsin, alcohol dehydrogenase, albumin, casein, and keratin) resulted in 704 proteins. Of these proteins, 425 (FOCC IDs) were also identified in the SG tissue proteome **(Supplementary File 1: Table S6**), and as such, we consider these with high confidence to be SG-secreted saliva proteins. A comparative analysis determined that 20%, 17%, and 6.8% of these 425 saliva proteins were identified in 1) the suite of SG proteins with in silico prediction of secretion in the present study (185), 2) a published SG transcriptome (Stafford-banks et al 2014), and 3) a published analysis and annotation of SG-enriched transcript sequences (Rotenberg et al., 2020), respectively. Of the 29 saliva proteins identified as SG-enriched transcript sequences of *F. occidentalis* (Rotenberg et al., 2020) (**Supplementary File 1: Table S7**), 19 (65.5%) were upregulated in SG tissues of females compared to males, and none were apparently influenced by infection.

Most of the SG-secreted saliva proteins (97%) had putative blastx descriptions of known proteins, while the remaining (13 proteins) are uncharacterized. Such proteins included three (SG5, SG20, SG40) previously determined to be transcript-enriched in SG tissues relative to whole bodies (Rotenberg et al 2020). The proportion of secreted saliva proteins assigned GO annotations for the four sex-treatment cohorts was 24% (M-NV), 26.5% (M-V), 28.3% (F-NV), and 26.9% (F-V). Predictions of subcellular localization patterns placed the secreted saliva proteins in diverse cellular compartments, with the majority localized to the cytoplasm, extracellular matrix, and mitochondria. GO terms overrepresented in the saliva protein dataset were ‘peptidase activity’, ‘carbohydrate metabolic process’, ‘hydrolase activity (acting on glycosyl bonds)’, and ‘ribosome’ (**Table 1; Supplementary File 1: Table S8**).

**Table 1:** ^1^Enriched gene ontology (GO) terms associated with proteins identified in secreted saliva relative to the salivary gland proteome of *Frankliniella occidentalis* adults. A complete list of saliva proteins in each GO class (MF = molecular function; BP = biological process; CC = cellular component) is provided in Supplementary File 1: Table S8.

| GO ID | GO Name | Class | FDR | p-value |
| --- | --- | --- | --- | --- |
| GO:0140096 | Catalytic Activity, Acting On A Protein (L3)<sup>2</sup> | MF | 1.02E-10 | 1.26E-13 |
| GO:0008233 | Peptidase Activity (L4) | MF | 2.03E-10 | 5.04E-13 |
| GO:0016787 | Hydrolase Activity (L4) | MF | 5.51E-10 | 2.05E-12 |
| GO:0009056 | Catabolic Process (L3) | BP | 1.21E-05 | 6.09E-08 |
| GO:0003824 | Catalytic Activity (L2) | MF | 1.21E-05 | 7.49E-08 |
| GO:0005975 | Carbohydrate Metabolic Process (L4) | BP | 1.40E-04 | 1.22E-06 |
| GO:0016798 | Hydrolase Activity, Acting On Glycosyl Bonds (L4) | MF | 1.40E-04 | 1.18E-06 |
| GO:0044281 | Small Molecule Metabolic Process (L3) | BP | 0.001 | 1.11E-05 |
| GO:0006091 | Generation Of Precursor Metabolites And Energy (L4) | BP | 0.015 | 1.63E-04 |
| GO:0003735 | Structural Constituent Of Ribosome (L3) | MF | 0.015 | 1.89E-04 |
| GO:0016829 | Lyase Activity (L3) | MF | 0.024 | 3.23E-04 |
| GO:0005829 | Cytosol (L3) | CC | 0.043 | 6.91E-04 |
| GO:0005840 | Ribosome (L6) | CC | 0.043 | 6.87E-04 |

<sup>1</sup>OmicsBox v2.1.14 (Götz et al., 2008) was used to perform the GO enrichment analysis using the two-tailed Fisher's exact test with a false discovery rate (FDR) of 0.05.
<sup>2</sup>Numbers in the parentheses (L#) indicate the hierarchical level of the GO term

### 3.8. TSWV-infection of *F. occidentalis* diversified putative processes of the male saliva secretome

Distribution of proteins into their most specific GO terms (multi-level GO analysis, terminal nodes) for each sex-virus treatment combination revealed that infection influenced the composition (type of proteins) and diversity of provisional biological processes associated with the saliva secretome (S**upplementary Fig. S4**). The virus effect was most notable for saliva of V males compared to NV males (**Supplementary Fig. S4.B vs S4.A**). While saliva of both male treatments contained proteins indicating carbohydrate metabolic and catabolic activities, there was an apparent shift from proteins overrepresented in ‘translation’ (particular subset of 40s and 60s ribosomal proteins, 20%), ‘small molecule metabolic process’, ‘cellular component organization or biogenesis’, to ‘biosynthetic process’, ‘protein metabolic process’ (amide and nitrogenous compounds), ‘cellular nitrogen compound and amino acid metabolic processes’. The virus-associated effect resulted in male saliva processes that closely mirrored those associated with NV females (**Supplementary Fig. S4.B vs S4.G**). In contrast, saliva secretomes of V females were less functionally diverse than NV females, whereby the most apparent modification was the overall reduction in proteins inferring “Hydrolase activity, acting on glycosyl bonds,’’ (**Supplementary Fig. S4.I vs S4.J**) including lysozyme, myrosinase, β-galactosidase, α-glucosidase, chitinase, N-acetylglucosaminidase, glycoside hydrolase 5, and lysosomal α-mannosidase. These proteins are diversified into different GO groups in infected saliva with ‘Hydrolase’ as the predominant category indicating potential role in other metabolic processes under infection status. Common ontologies across all four sex-treatment combinations were “peptidase activity” (primarily proteases - thus an indication of protein degradation), “oxidoreductase activity”, and “ion binding” with a 40% overlap in sequence IDs of these GO-annotated sequences. However, within these three GO categories, infected males contained proteins not identified in the other three treatments, notably members of multiple families of proteasome subunits.

### 3.9. Saliva-secreted proteins unique to TSWV-infected adults indicate energy generation, consumption and protein turnover

Identities (Focc OGS IDs) of the 425 secreted saliva proteins were used to determine shared and unique sets of proteins across the four sex-treatment combinations. A common core set of proteins (42%) were secreted in the saliva regardless of sex and virus infection, while 10.1% were shared exclusively between infected males and females (**Fig. 5A; Supplementary File 1: Table S9)**. There were proteins identified only in non-exposed males (5.2%; 62% were ribosomal proteins) or females (1.4%), and virus infection appeared to increase the occurrence of saliva proteins (i.e., number of different proteins) unique to males (11.4%) and females (6.8%). *In silico* analysis of putative, physical PPI networks (STRING) predicted for the 425 saliva proteins (84% mapped to the *D. melanogaster* protein database) and the subsets uniquely found in saliva of infected cohorts (**Fig. 5A**: M-V (49), F-V (29) and M-V/F-V (43)) revealed several evolutionarily conserved clusters of physically-interacting proteins in networks that further distinguished the sexes under infection (**Fig. 4B & Fig. 5B**). Unique to infected females, there was a predicted PPI cluster of seven proteins associated with the ATP-generating and amino acid-producing intermediates of the tricarboxylic acid pathway, the conserved energy-producing process of the mitochondrion. Unique to infected males, one PPI cluster of six proteasome subunits of the 20S catalytic core involved in proteolysis [Prosbeta 1, 4, 6, and 7; alpha 5 and 6] was predicted; prosbeta 1 and 4 have been implicated in cellular response to DNA damage. Other PPI clusters contained proteins shared and unique to infected males or females. The most prominent was a PPI cluster of 18 proteins associated with actin microfilament organization and movement, the extracellular matrix, laminin-interacting proteins of basement membranes, and muscle fibers. Integrinβ-PS (mysopheroid isoform C in *D. melanogaster*, laminin receptor) was central to that cluster (**Fig. 4B**). Smaller PPI clusters indicating aldehyde catabolism/detoxification and fatty acid degradation were also apparent.

**Figure 5.**
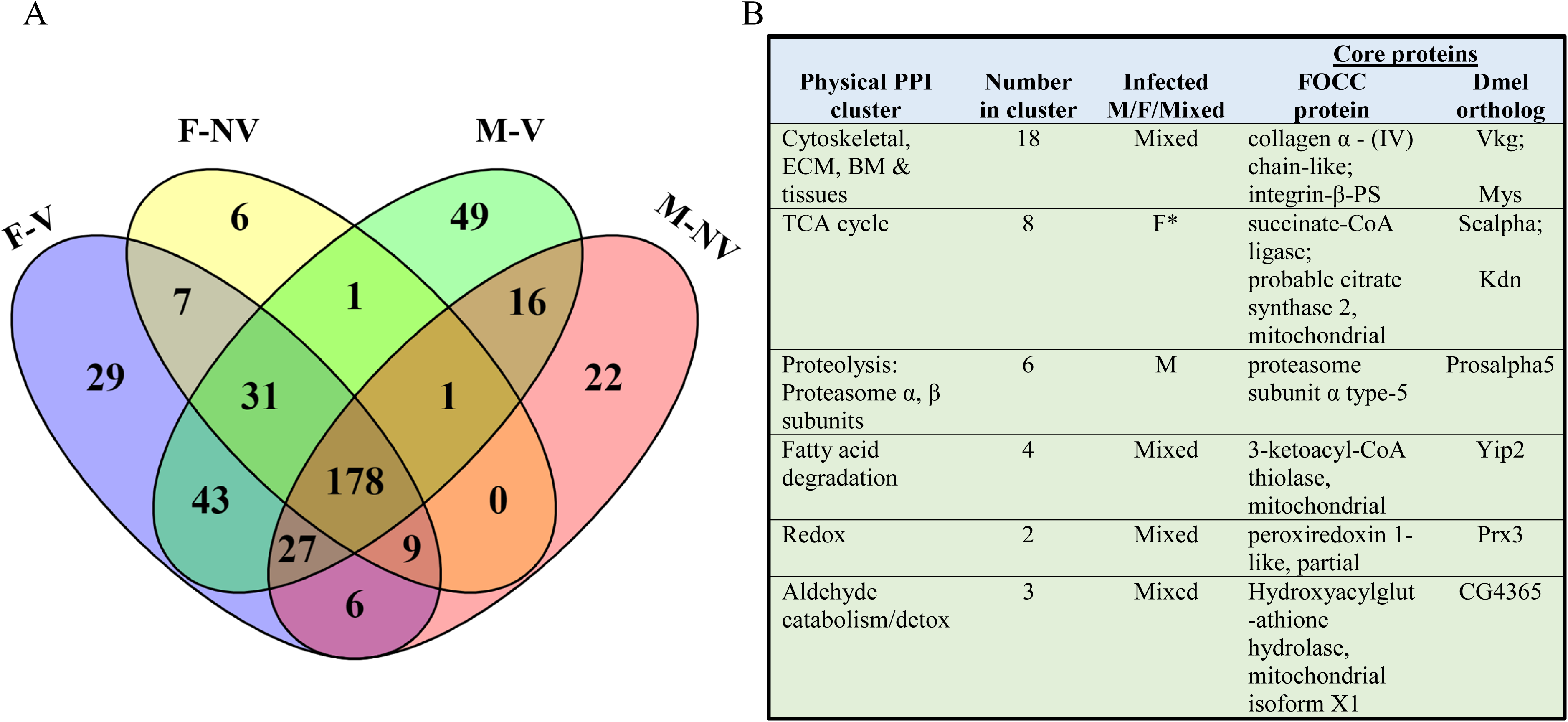
Unique and shared saliva protein sequences identified from tomato spotted wilt virus-infected males and females of *Frankliniella occidentalis*. (A) VENN diagram depicting overlaps between the four sex-treatment cohorts: non-exposed males (M-NV), non-exposed females (F-NV), infected males (M-V) and infected females (F-V). Each cohort represented pools of males or females generated from 12 independent TSWV-acquisition assays (larval thrips) on infected plant tissue; in parallel, each assay included adult cohorts generated from larval cohorts fed on healthy plant tissue (mock-treatment). (B) Summary of predicted protein-protein interaction (PPI) networks (clusters of physically-interacting proteins) among saliva proteins unique to TSWV-infected adults. Refer to Figure 4B for illustration of protein clusters, core interactors, and network annotations of the saliva PPI networks.

### 3.10. Overlap between SG vDAPs and saliva of infected adults

Of the collective 78 vDAPs in SGs of *F. occidentalis* adults, 10 were identified in the secreted saliva of infected adults - three and seven proteins in males and females, respectively. Specifically, the DAPs integrinβ-PS and serine protease 29 (both upregulated) and peroxiredoxin-4 (downregulated, the only DAP overlap with males) in infected females were identified in the saliva. In infected males, three upregulated SG proteins - an uncharacterized protein, hypothetical protein, and nidogen - and four downregulated proteins - pro-dipeptidase, peroxiredoxin 1 and 4, and glycerol-3-phosphate phosphatase were secreted in the saliva.

### 3.11. Ancient lateral gene transfers (LGTs) expressed in *F. occidentalis* salivary glands and secretome

As part of the annotation of the *F. occidentalis* genome, two LGT events of different bacterial origins, each exhibiting gene expansions and showing evidence of purifying selection and expression (transcript) in whole bodies of *F. occidentalis*, were described (Rotenberg et al., 2020, refer to Additional File 1). These two LGTs (i.e., mannanase and levanase) are glycoside hydrolases (GH), members of a class of proteins involved in carbohydrate metabolism (Drula et al., 2022; Cazy database: https://doi.org/10.1093/nar/gkab1045. In the present study, one of the three paralogs of the mannanase gene (OGS ID: FOCC012782, Genbank acc: XP_026285291.1) was identified in the male and female SG proteomes (**Supplementary File 1: Table S3**), and the same mannanase and one of the copies of the levanase genes (OGS ID: FOCC006419, Genbank acc: XP_026287851.1) were identified in SG-secreted saliva (**Supplementary File 1: Table S6**). The MSStats analysis of the SG proteome revealed that the mannanase (LGT) protein was less abundant in M-NV compared to F-NV (log2 fold change = −0.44, *P* = 00742), and this differential sex/virus effect was supported by a subtle, but significant sex-virus treatment interaction (*P* = 0.0407) in a simple two-factor ANOVA (JMP pro v15, SAS Institute) on normalized abundance values (log2-transformed). The levanase (LGT) protein identified in the saliva of both sexes was filtered from the SG proteome because only one unique peptide identified the protein (criteria = at least 2 unique peptides).

### 3.12. Putative orthologs of *F. occidentalis* saliva proteins in other piercing-sucking herbivorous arthropods

A comparison between the *F. occidentalis* secreted saliva protein sequence dataset (425) with published saliva protein datasets of three representative hemipterans (aphid, planthopper, psyllid) and one arachnid (two-spotted spider mite) showed that 26% (111) of the *F. occidentalis* saliva proteins (both sexes regardless of infection status) were provisional orthologs of saliva proteins in at least one of the four queried arthropod species (**Supplementary File 1: Table S10**). Approximately 4% of the thrips saliva proteins identified putative orthologs in at least two of the queried hemipteran species datasets. The rank of arthropods in order of the most to least number of matches with *F. occidentalis* saliva proteins was *D. citri* (the Asian citrus psyllid) (18.1%), *B. tabaci* (9.2%) *N. lugens* (small brown planthopper) (6.3%), *T. urticae* (two-spotted spider mite) (5.6%), *M. euphorbiae* (the potato aphid) (2.8%), and *M. persicae* (the green peach aphid) (2.8%). The ranking may, in part, reflect the size of the subject sequence databases for the comparative species (refer to Section 2.9.). Nonetheless, there were some identifiable trends. The similarity between *D. citri* and *F. occidentalis* was dominated by the presence of several secreted saliva proteins associated with cell structure, protein synthesis and maintenance. Notable groups include several cytoskeletal and extracellular matrix proteins (actin, myosin, papilin, lamilin, tubulin, titin, and basement membrane heparin proteoglycan), 15 ribosomal proteins, and four heat shock proteins. Proteins shared between *F. occidentalis* and *T. urticae* indicate enzymatic activities that potentially necessitate interactions with plant physical and chemical components (Guo et al., 2020; Kettles and Kaloshian, 2016; Ray and Casteel, 2022; Sharma et al., 2014; Subramanyam et al., 2021), including catabolic enzymes such as carbohydrate degrading enzymes (β-galactosidase, N-acetylgalactosaminidase, chitinase), protein degrading enzymes (cathepsin-L&B, serine proteases) and detoxification enzymes (glutathione-s-transferase, superoxide dismutase, carbonyl reductase). The majority of the proteins (11) shared between *F. occidentalis* and *B. tabaci* were enzymes that may interact putatively with plant components. Whiteflies and aphids are phloem feeders, and very few (2.8%) aphid saliva proteins overlap with the thrips saliva proteins, indicating a vast difference in the feeding physiology of these two groups. This difference could be attributed to more than the technical approaches to study the saliva proteome of the respective insect members.

### 3.13. Other secreted proteins of *F. occidentalis*

Of the 279 proteins identified in the feeding solutions but not detected in the SG tissue proteomes (**Supplementary File 1: Table S11**), 24% and 23% were female- and male-specific, respectively. Regardless of sex, most of these have provisional annotations (GOs) associated with amide and carboxylic acid metabolic process (6%), intrinsic component of membrane (19%), nucleotide binding (30%), and serine-type endopeptidase (12%). Some of the protein identities in this group include hypothetical proteins (25), myrosinase (22), uncharacterized proteins (11), trypsin (10), and juvenile hormone (8). The origins of these proteins remain to be determined.

### 3.14. TSWV proteins were identified in *F. occidentalis* salivary glands and secreted saliva

TSWV glycoprotein precursor, nucleocapsid protein (N) and non-structural protein NSs were identified in both the SGs and saliva of the infected adults. The glycoprotein precursor had four unique peptides with a single peptide aligned to the G_N_ region and three peptides aligned to the G_C_ region in SGs of infected females only. Nucleoprotein (N) and NSs proteins were identified with 16 and eight peptides, respectively, across SG samples of TSWV exposed adults.

The normalized abundance of NSs protein was significantly greater in SGs of infected males compared to infected females (**Figure 6**, *P* = 0.032, Pooled T-test). Reflecting the infection status of thrips, secreted saliva of TSWV-infected females contained all three proteins (Glycoprotein precursor, N, and NSs), whereas saliva of infected males contained glycoprotein precursor and N proteins. Saliva of infected males contained nine and six unique peptides for glycoprotein precursor and N protein respectively, whereas saliva of infected females contained three, four, and two unique peptides for glycoprotein precursor, N, and NSs proteins. The RNA-dependent RNA polymerase (L, a viral structural protein known to in low abundance in the virion, (Mohamed et al., 1973)) and movement protein (NSm, a non-structural protein known for its role in plant cell-cell movement, Lewandowski and Adkins, 2005) of TSWV were not detected, which may indicate these proteins were below detection level.

**Figure 6:**
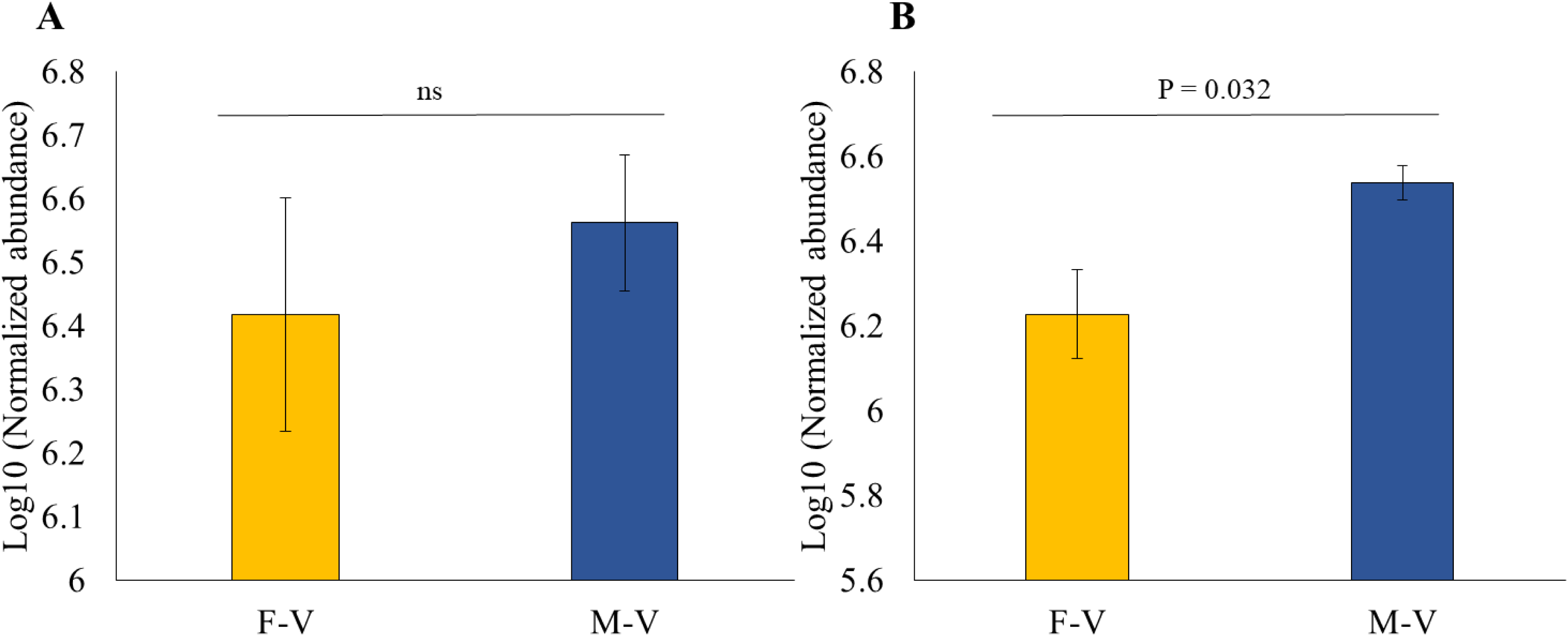
Tomato spotted wilt virus proteins expressed and differentially-abundant in *Frankliniella occidentalis* salivary glands with respect to sex. Log_10_-transformed normalized abundance of (A) TSWV nucleocapsid (N) protein and (B) non-structural protein NSs. Bars represent the mean ± standard error of the mean (n = four independent experiments). Pooled t-tests were performed to determine statistical differences (*P* < 0.05) between sexes.

All measures were taken to prevent contamination of samples from the time of preparing virus inocula, to preparing the treatment groups of thrips (NV and V), to rearing the treatments to adulthood, to dissecting glands and collecting saliva, and to delivering samples to the proteomics facility. Nonetheless, peptides that indicated TSWV proteins (N, NSs, glycoprotein) were detected in the salivary gland samples of the non-exposed (NV) treatment, albeit to a significantly lower abundance than that quantified for SGs of the virus-exposure treatment (V; 3.7-fold and 2-fold higher for N and NSs, respectively). In addition, the presence of specific peptides that matched TSWV proteins indicated a more sporadic, patchy occurrence across the biological replicates of the NV samples (male and female) compared to the V samples (**Supplemental Figure S4**). Searching NCBI protein databases, published thrips viromes (Chiapello et al 2021; Thekke-veetil et al 2020) and genome scaffolds of *F. occidentalis* did not recover sequences 100% identical to the TSWV peptides detected in the SG samples. Collectively, our diagnostic assessment of viral proteins in SG samples was most likely due to superficial contamination (e.g., tissue harvesting or post-processing steps) and not infection of glands in the NV treatment. No TSWV peptides were detected in saliva collected from NV treatment groups.

## 4. DISCUSSION

Our study provides the first description of the SG proteome and secretome (saliva) of a thysanopteran, and in the present case, a herbivorous pest and vector of plant-pathogenic viruses that circulate, replicate and persist over the lifetime of the insect. In line with other arthropod pests (Sharma et al., 2014), salivary glands of *F. occidentalis* adults expressed a common core of proteins with diverse physiological functions, including carbohydrate metabolism, catabolism, proteolytic activities, oxidation-reduction of diverse substrates, and ion binding. The most abundant proteins documented for SGs of male and female thrips, and commonly abundant in other invertebrate tissues, were lipoproteins (lipophorin precursor, apolipophorin and vitellogenin). Lipoproteins comprise a large superfamily of proteins (Babin et al., 1999) that, in arthropods, are expressed in fat bodies, secreted into the hemolymph, and are involved in intra-and extracellular transport of lipids (e.gs., fatty acids and sterols) to various tissues during development. In some cases these proteins have been co-opted for other functional roles in herbivorous vectors. For example, vitellogenin, known for its role in egg production, has evolved functional roles in facilitating host feeding by *Laodelphax striatellus* (the small brown planthopper) by attenuating plant defenses (Ji et al., 2021), and has emerged in insect saliva proteomes, including plant pests (Huang et al, 2020; Wu et al, 2021). Ultrastructure examination of *F. occidentalis* SGs of adults showed aggregates of cells containing numerous lipid droplets (Ullman et al., 1989), which are ER-originating storage organelles that serve to store lipids, primarily triacylglycerides, and to regulate energy homeostasis in eukaryotes, including insects (reviewed in (Kühnlein, 2012)). Two perilipin-4-like proteins were identified in the SG proteome of both *F. occidentalis* sexes, regardless of virus infection, and one was identified in the saliva. Perilipin family members are lipid droplet-associated proteins involved in lipid mobilization and lipid accumulation in mammals (Olzmann and Carvalho, 2019) and other metazoans, including *D. melanogaster* (Kühnlein, 2012). The transcripts that encode the perilipin-4-like proteins identified in this present study were originally identified as SG tissue-enriched transcripts ranging between 2.5 to 25-fold more abundant in SGs compared to whole bodies of adults (Rotenberg et al., 2020, refer to Additional File 2, Supplementary Table 11 of the citation). The roles of these putative lipid storage proteins in *F. occidentalis* with regards to lipid mobilization during times of starvation, storage during thrips development, or TSWV infection - lipid ‘hijacking’ is commonly ascribed to arboviruses in their mammalian and vector hosts (reviewed in (O’Neal et al., 2020)) - warrants investigation.

We documented a robust sex-biased SG proteome for *F. occidentalis* as evident by the large proportion of sDAPs between sexes (over 50%) and their provisional cellular roles, regardless of virus infection status. For those evolutionarily conserved proteins (i.e., GO evidence), gene ontologies indicated SG tissue-enrichment of active translational machinery and protein synthesis in females compared to males, perhaps underscoring the allocation of energy (via active feeding) to support the metabolic cost of egg production (Smykal and Raikhel, 2015). In contrast, the male SG proteome indicated diverse catalytic activities on cell and subcellular substrates, many of which infer fundamental, molecular functions. Sex-biased expression and adaptive innovations of SG proteomes have been documented for well-characterized blood-feeding arthropods (Ribeiro et al., 2018, 2016). However, while there are reports for SG proteome plasticity and adaptations due to changes in food sources of plant-feeding arthropods (Mathers et al., 2017), there are no comparable reports of sex-biased SG proteomes for these species, most likely, in part, due to the focus on thelytokous parthenogenic females (aphids) and juvenile stages. Males of *F. occidentalis* tend to spend less time feeding (Stafford-Banks et al., 2014) and have short life spans (Reitz, 2009) compared to females, and we speculate that sex-biased SG proteomes of *F. occidentalis* may underscore, in part, these sexually-dimorphic traits.

Salivary gland tissues and saliva play crucial roles in virus transmission to hosts. We hypothesized that TSWV infection modulates the SG proteome of *F. occidentalis*, and that the modulation is sex-biased. Our findings support this two-component hypothesis by the number, differential abundance, identities and generalized roles of the proteins influenced by virus infection. Our study documented little overlap in vDAPs between the sexes - only one protein was shared (peroxiredoxin-4-like), down-regulated in both. Peroxiredoxins participate in the detoxification of peroxides which cause oxidative damage to cells (Perkins et al., 2015) ; however, they have been implicated in H_2_O_2_-associated signaling and innate immunity (e.g., involved in PAMP signaling) (Rhee, 2016). An RNAseq analysis of TSWV-infected first instar larval bodies also revealed a downregulation of transcripts encoding this protein (Schneweis et al., 2017) (Scheneweis et al., 2017). Perhaps perturbation of peroxidase levels in SGs of adult *F. occidentalis* is a generalized response to TSWV infection, and down-regulation promotes a supportive environment for the virus. Male SGs exhibited a larger proteomic response to TSWV infection than female SGs. Moreover, inferred protein functions and predicted functional protein networks - known proteins that physically interact or crosstalk within or between pathways - associated with infected male SGs are consistent with a physiological state of reduced respiration, and thus energy production (mitochondrial shutdown), and the accumulation of fatty acids. In addition, there appeared to be an orchestrated modulation of nuclear and nucleolar activities associated with mRNA availability and ultimately expression as documented by the upregulation of members of the spliceosome pathway coupled with a downregulation of proteins associated ribosomal biogenesis, histone methylation, and mRNA unwinding. Together, these subcellular processes (mitochondrial and nuclear) perturbed by TSWV in males point to a further slow-down in metabolism under infection. Whether TSWV infection causes damage to SG tissue and/or alters the physiological fitness of males (eg. longevity) compared to non-exposed males or infected females has yet to be determined.

Several male vDAPs identified in this study have been reported in other host-virus pathosystems to play critical roles in viral processes. These virus processes include:

***i) Replication and infectivity:*** Spermidine/spermine acetyltransferase 1-like (SAT1-like) was determined to be more abundant in SGs of TSWV-infected males compared to non-exposed males. SAT1 is a key enzyme involved in the catabolism of polyamines, and its abundance is positively regulated by polyamine levels in the cell (reviewed in Firpo and Mounce, 2020). Polyamines, such as spermidine, spermine and putrescine, are aliphatic polycations produced by eukaryotic cells that bind nucleic acids, proteins, or lipids and are essential for healthy cell growth and longevity (Handa et al., 2018; Igarashi and Kashiwagi, 2019). Polyamines have also been shown to be indispensable with regards to replication of diverse families of animal-infecting viruses (Gibson and Roizman, 1971; Mastrodomenico et al., 2019; Mounce et al., 2016), including bunyaviruses. Rift Valley fever virus (RVFV) was functionally shown to require particular polyamines for replication and infectivity *in vitro*, and it was determined that spermidine associated directly and transmitted cell-to-cell with the virion, serving an essential role in infectivity (Mastrodomenico et al., 2020). Diverse plant viruses, including turnip yellow mosaic virus (TYMV) and Belladonna mosaic virus (BDMV) likewise incorporate host polyamines (spermidine or putrescine) in the virion (Balint and Cohen, 1985; Beer and Kosuge, 1970; Johnson and Markham, 1962; Virudachalam et al., 1983b), with some evidence suggesting a role in virion stability (Savithri et al., 1987; Virudachalam et al., 1983a). Given the positive regulation of SAT1 levels by polyamine levels in animal cells (Firpo and Mounce, 2020), the increased abundance of the SAT-1-like protein in TSWV-infected male SG cells may indicate, in part, an increased catabolism of polyamines that support TSWV infectivity, replication, and/or cell-cell movement.
***ii) Endocytosis (entry) and vesicle-mediated transport:*** Sorting nexin-9, a putative ortholog to mammalian LST-4 sorting nexin - a protein that modulates endocytosis and endosomal sorting (Cullen, 2008) - was more abundant in TSWV-infected male SGs compared to non-exposed males. Animal-infecting bunyaviruses enter cells via receptor-mediated endocytosis (Guardado-Calvo and Rey, 2017), and by analogy, it has been proposed that TSWV may enter thrips vector cells by the endocytic pathway (Chen et al., 2019; Whitfield et al., 2005). Insights from other animal-infecting (alphavirus, Sindbis virus) and plant-infecting (begomovirus, Tomato yellow leaf curl virus) arboviruses demonstrated the role of host sorting nexins in virus replication and intracellular trafficking in vector cells, respectively (Schuchman et al., 2018; Xia et al., 2018). In TSWV-infected males, in relation to non-exposed males, SG abundance of ADP ribosylation factor-like protein 8B/A (ARF8), a GTP-binding protein reported to be a lysosomal membrane protein and implicated in lysosomal trafficking (Khatter et al., 2015), was determined to be in lower abundance. Regulation of lysosomal trafficking may be an indication of a perturbation of the late stage endosomal transport of cargo to the lysosome (i.e., possible endolysosomal late-entry of viruses; (Lozach et al., 2010)) and/or protein degradation. Silencing of a different ARF (ARF1) in *Psammotettix alienus* (European grass feeding leafhopper), resulted in significant reduction in virus accumulation and dissemination of wheat dwarf virus to the hemolymph of the hopper vector (Wang et al., 2019).
***iii) Intracellular and cell-cell movement:*** Protein 4.1 homolog (3-fold UP in MV; Dmel cora), a cytoskeletal protein that interacts directly with other cytoskeletal proteins (Baines et al., 2014) to form a scaffold with membranes, has been shown to be involved in virus entry and/or trafficking. For example, protein 4.1 homolog was determined to be crucial for Zika virus (mosquito vector) entry into erythrocytes (Su et al., 2021). In the present study, *F. occidentalis* protein 4.1 homolog was predicted to functionally interact with an actin-associated spectrin alpha chain protein (UP in MV) and integrin beta-PS (UP in FV), structural proteins involved in cell-cell interactions (i.e., cytoskeletal structure and endo/epithelial integrity/cell signaling, respectively). Many arboviruses use the cytoskeletal network of their insect host or vector to complete their life cycles (reviewed in Khorramnejad et al., 2021). We postulate that cytoskeletal vDAPs in *F. occidentalis* may enable TSWV movement within and between SG cells.

Generalized biological roles inferred from vDAPs in female SGs highlighted several up-regulated proteins associated with tissue integrity. Integrin-beta-PS (Dmel myospheroid, isoform c), cell-surface receptor of laminin (BM protein), is a conserved protein that contributes to endodermal integrity and cell-to-cell adhesion (eg., intestinal stem cell and surrounding visceral muscle cells in adult Drosophila midguts) (Lin et al., 2013). In the present study, the thrips integrin-b-PS (UP in F-V vs F-NV) was predicted to be at the core of the cell-cell interaction protein network (cytoskeleton/BM/ECM) of male infected saliva, possibly indicating dysregulation of these proteins in males and female SGs in response to TSWV. β integrins have been demonstrated to be receptors for some viruses in the family *Hantaviridae* (Gavrilovskaya et al., 1999, 1998; Matthys et al., 2010). Because integrin-b-PS has a known tissue distribution in insects that is relevant to the TSWV dissemination pathway and the role of related proteins in entry of other viruses in this order, these proteins are of particular importance for functional analysis of their role in virus entry and movement within the thrips vector. Another protein, flightin, which is ubiquitously present in many arthropods is an integral component of muscle structure of arthropods (Chakravorty et al., 2017; Xue et al., 2013), most notably associated with wing muscles. A putative cuticle protein CPG38 (elastin-like) was one of the most abundant vDAPs in female SGs. A previous study identified two cuticular proteins (CP-V and endoGN) expressed in *F. occidentalis* larval guts and that interacted directly with TSWV G_N_, the viral attachment protein (Badillo vargas et al., 2019) and CPs have been implicated in other plant virus-vector interaction studies (Deshoux et al., 2018). Functional characterization of these noted male and female vDAPs warrants further study to definitively determine their putative roles in TSWV-thrips interactions in SG tissues.

Our study exemplifies the necessity to directly identify components of SG-secreted saliva in order to resolve potential virus effects on the vector-plant interface. Our findings documented the effect of TSWV infection on cellular and subcellular enzymatic functions of SG cells - nuclear, mitochondrial, intracellular cytoskeletal network, membrane biochemistry - and little to no effect on plant cell wall-degrading enzymes or proteins associated with detoxification of plant chemistries. On the contrary, the core set of proteins identified in the saliva of *F. occidentalis* (regardless of virus infection status) included carbohydrate- and protein-degrading enzymes that have also been found in other insects within the similar feeding guild (Jonckheere et al., 2016; Lomate and Bonning, 2016). We determined that most of the thrips saliva proteins did not contain predicted signatures of secretion. This finding may be explained by the presence of non-canonical secretion signals, errors in predictions, or proteins that hitchhike a coordinated active mechanism of transporting cargo across plasma membranes (e.g., exocytosis Nawaz et al., 2020). Conversely, only 20% of SG proteins predicted to be secreted were detected in the saliva - perhaps those unaccounted proteins were below the level of detection, technical limitations precluded detection of post-translational modifications, or SG-secreted proteins may be destined for the ECM and not the SG epithelium.

The limitations of working with small insects (< 1 mm body length) constrained the saliva to a qualitative analysis of pooled samples across multiple (12) independent repetitions of the underlying experiment. Nonetheless, our study provided evidence of virus effect based solely on the occurrence (presence or absence) of identified saliva proteins. Saliva secreted by TSWV-infected adults of *F. occidentalis* in this study contained putative homologs of proteins in hemipteran vectors that have been functionally characterized to interact with plant defense pathways. For example, two calcium-binding proteins, regucalcin and calmodulin, were reported to inhibit calcium-derived callose formation in the phloem during aphid feeding (Reviewed by van Bel and Will, 2016). An odorant-binding protein (OBP11) identified in saliva of *Nilaparvata lugens* (brown planthopper), was shown to promote feeding by suppressing the production of salicylic acid - a phytohormone that regulates plant defense against pathogens - in rice plants (Liu et al., 2021). Unique to TSWV-infected females, ATP-generating and amino acid-producing intermediates of the tricarboxylic acid pathway, evidence of active energy generation, were enriched in the saliva. Unique to TSWV-infected males, two carbohydrate-degrading enzymes, β-galactosidase and glycoside hydrolase family 32 (GH32), were identified in the saliva. Furthermore, two glycoside hydrolases (mannanase and levanase) previously reported to be two ancient bacterial LGTs in the *F. occidentalis* genome (Rotenberg et al., 2020) were identified in saliva of both sexes, regardless of virus infection. These two thrips GHs likely play roles in plant cell-wall degradation and feeding (carbohydrate metabolism), as documented for other crop pests (Subramanyam et al., 2021).

The sex-biased effect of the virus on SG physiology of *F. occidentalis* extended to the saliva. Virus infection diversified the ontologies of saliva proteins secreted by males, and patterns indicated a shift from a preponderance of translational machinery (ribosomal subunits) to protein metabolism (proteasome subunits, peptidases, digestive serine proteinase). These shifts in saliva composition under infection may contribute, in part, to enhanced host probing and feeding patterns exhibited by TSWV-infected males of *F. occidentali*s (Stafford et al., 2011). Furthermore, the diversification of male saliva functions mirrored those inferred from non-exposed female thrips, supporting the hypothesis that infected males exhibit similar feeding behaviors as non-infected females (Stafford et al., 2011). The virus effect on putative female saliva functions was more subtle, a finding that reflects the subtle effect of virus on the glands.

Proteins known for their roles in intracellular cytoskeletal networks, BMs, and the ECM, i.e., proteins associated with cell adhesion, integrity and migration, were also deposited in the SG-secreted saliva of *F. occidentalis*. Our comparative analysis identified putative orthologs of these cytoskeletal and structural membrane proteins in saliva of plant-feeding hemipterans (aphids, phyllid, planthopper), an indication of a common occurrence in piercing-sucking insects. The delivery mechanism for these structural proteins into saliva of plant-feeding insects is unknown. In the present study, these structural proteins (cellular and tissue) were overrepresented in the saliva of TSWV-infected adult thrips, and together, these proteins formed the largest predicted functional interaction network (including integrin, laminin, filamin, vinculin, nidogen, collagen, calmodulin and others). Here we propose various scenarios (hypotheses) for the occurrence of these types of protein in the *F. occidentalis* saliva. One, proteins perturbed by TSWV infection might be exocytosed along with virions or viral proteins (Chen et al., 2021; Nawaz et al., 2020). Similar to other persistent plant viruses, TSWV is proposed to be transported via an intracellular path and exocytosed into the salivary lumen (Chen and Wei, 2020; Ullman et al., 1995). Two, the proteins may interact directly with TSWV proteins and be subsequently packed into virions released into the salivary duct. For example, a sorting nexin (different from the sorting nexin identified in this study) is also known to be packed into the viral particles of mosquito-vectored arbovirus (Schuchman et al., 2018). Three, increased occurrence of cytoskeletal and tissue structural proteins in saliva may indicate signs of SG cell turnover to counter damage caused by TSWV accumulation in the tissues. The phenomenon of SG cell turnover is well-studied in vertebrates (reviewed in Aure et al., 2015), and it has been suggested that occurrence of cytoskeletal proteins in SGs of vectors of animal pathogens is associated with damage caused by infection (Sim et al., 2012; Telleria et al., 2014). An earlier study by Ullman et al (1989) documented the presence of autophagic vacuoles in *F. occidentalis* SG upon TSWV infection. These autophagic vacuoles may be a sign of histopathology, and the scenarios presented above warrant further ultrastructural microscopic investigations to elucidate export of cellular components and virions into the thrips saliva.

One novelty of our study was evidence of particular TSWV proteins in secreted saliva of infected *F. occidentalis adults*. TSWV NSs protein was identified in the saliva of infected females. TSWV NSs is a multifunctional viral effector protein that targets plant defenses against TSWV and thrips. NSs is a suppressor of local and systemic silencing in plants and insects thus protecting TSWV from this powerful antiviral defense response (Margaria et al., 2014; Takeda et al., 2002). NSs binds to long and short dsRNAs thereby preventing cleavage of viral RNA by dicer and loading of siRNAs into the RISC complex (Schnettler et al., 2010). The presence of this non-structural protein in the saliva indicates that this protein is co-delivered with viral particles into the plant cell and may be able to protect the virus from antiviral responses before the replication cycle begins in the host cell. NSs also inhibits jasmonic acid pathway mediated terpene biosynthesis by binding to MYC family transcription factors to increase the attractiveness and establishment of the *F. occidentalis* population on the host plant (Wu et al., 2019). Our study provides evidence of the delivery of this crucial viral protein to the host plants via thrips saliva, increasing the potential for co-delivery of a viral effector with inoculation of TSWV in nature. Furthermore, it remains for future studies to determine if co-delivery of NSs and particular thrips secreted saliva proteins identified in this study concomitantly attenuate plant defenses against pest and pathogen.

## AUTHOR CONTRIBUTIONS

**DR, DU, JBB, SPR** conceptualized the project; **DR** supervised the project; **DR** and **SPR** developed methodology; **SPR** investigated the project; **SPR** and **DR** formally analyzed the data; **DU, JBB, DR, SPR, SBM** discussed and interpreted findings; **SPR** and **DR** wrote the manuscript; **DR** and **SPR** designed and prepared visualizations. **AEW, SBM, DU, JBB, DR,** and **SPR** reviewed, provided insights and edited the manuscript.

## ACKNOWLEDGEMENTS

We thank Dr. Erik Soderblom, Dr. Arthur Mosley, Tricia Hoe, and Dr. Greg Waitt of the Duke Center for Genomic and Computational Biology for critical discussions of proteomics methodologies and parameters, and Drs. Jinlong Han and Kirsten Lahre for assistance with virus acquisition assays. This work is supported by NSF-NIFA Plant Biotic Interactions Program [grant no. 2018-67013-28495] from the USDA National Institute of Food and Agriculture.

## APPENDIX. Supplementary Materials

The following are the Supplementary Tables and Figures that support this manuscript:

Supplementary File 1: Workbook of large supplementary data tables, including salivary gland raw peptide sequences and metrics

Supplementary File 2: Raw peptide sequences and metrics for saliva-secreted and other proteins

Supplementary File 3: Proteins sequences of in-house clones of TSWV used in this study.

Supplementary Figure S1: Flowchart of approaches, downstream analyses and associated data visualizations

Supplementary Figure S2: Sample clustering (PCA), distributions (Boxplots) and most abundant salivary gland proteins (normalized protein abundance)

Supplementary Figure S3: Volcano plots of log_2_fold-change of normalized protein abundance between sexes

Supplementary Figure S4: Distribution of annotated proteins in secreted saliva by gene-ontology (GO) terms (pie charts)

Supplementary Figure S5: Occurrence and raw peptide abundance of TSWV proteins

