## Supplementary Figure S1 for "Sex-biased proteomic response to tomato spotted wilt virus infection of the salivary glands of *Frankliniella occidentalis,* the western flower thrips"

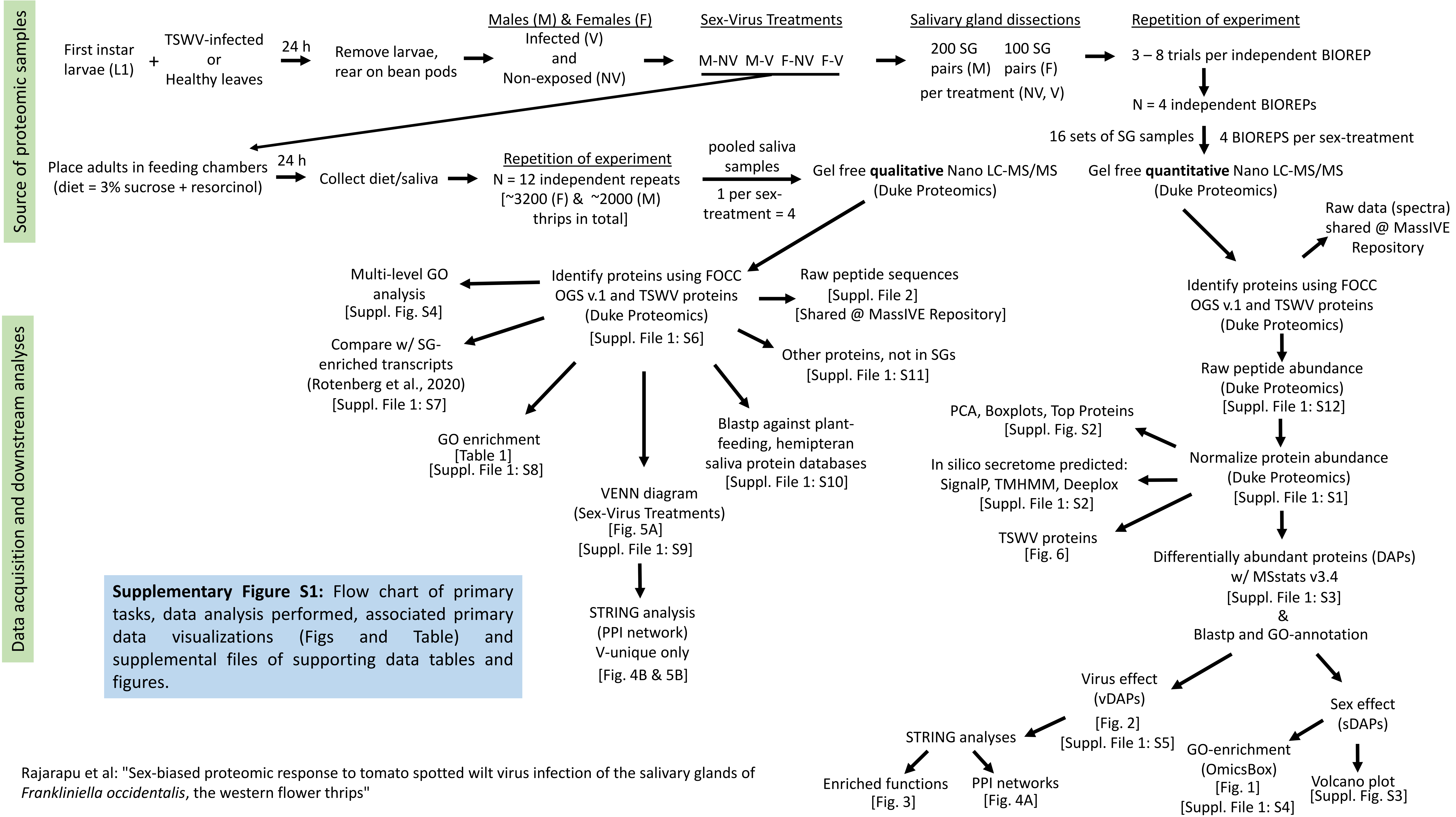


**Supplementary Figure S1:** Flow chart of primary tasks, data analysis performed, associated primary data visualizations (Figs and Table) and supplemental files of supporting data tables and figures.

Rajarapu et al: "Sex-biased proteomic response to tomato spotted wilt virus infection of the salivary glands of *Frankliniella occidentalis*, the western flower thrips"
