## Supplementary Figure S3 for "Sex-biased proteomic response to tomato spotted wilt virus infection of the salivary glands of *Frankliniella occidentalis,* the western flower thrips"

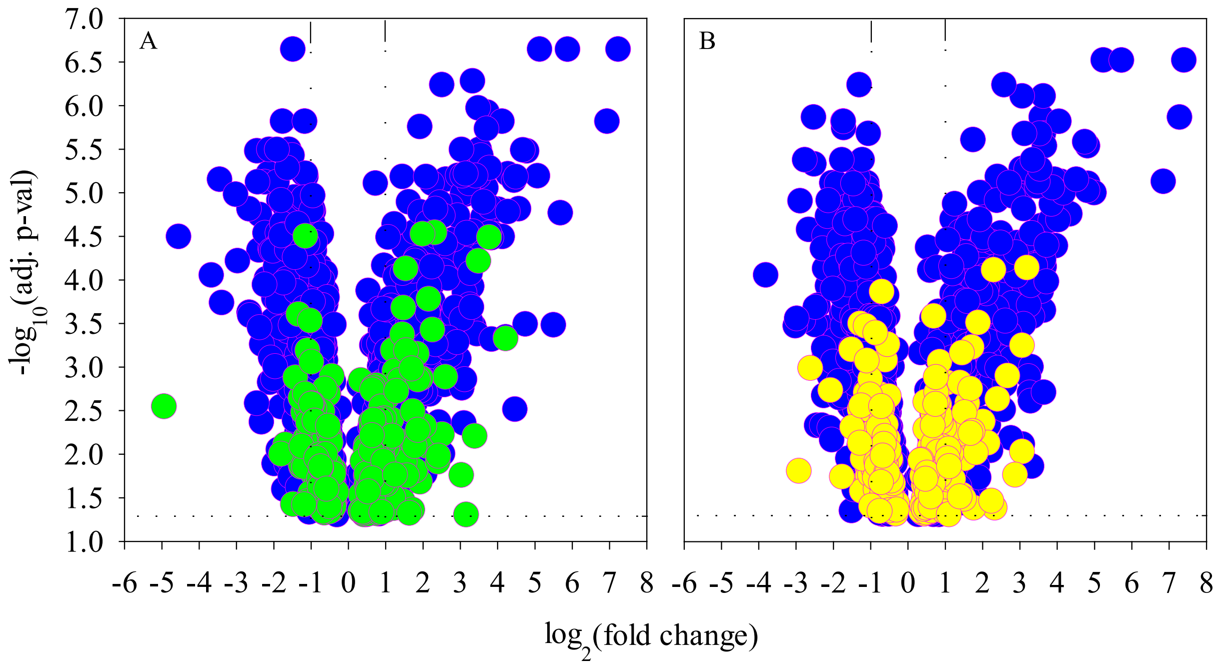


**Supplementary Figure S3**. Volcano plots depicting magnitudes of change in protein normalized abundance ($log_2$ fold change) of annotated proteins and associated levels of significance (adjusted p-value) for pairwise comparisons between adult female and male salivary gland tissues of *Frankliniella occidentalis*. A) non-exposed treatment (NV); B) tomato spotted wilt virus (TSWV)-exposed treatment (V). Blue symbols indicate differentially-abundant proteins modulated by sex (sDAPs) regardless of treatment (930 proteins); green and yellow symbols indicate sDAPs unique to the NV (245 proteins) and V (268 proteins) treatments, respectively. The horizontal dashed line indicates the cut-off for statistical significance (0.05), and vertical dashed lines demarcate sDAPs with fold change values of 2 (positive values = up-regulated in males in relation to females; negative values = up-regulated in females in relation to males). NV and V cohorts were generated by feeding first instar larvae on healthy, non-infected plant tissue or symptomatic, TSWV-infected plant tissue for 24 hours, respectively, followed by rearing to adulthood on green bean pods. Each point (symbol) represents the mean of N = 4 independent biological replications.

Rajarapu et al: "Sex-biased proteomic response to tomato spotted wilt virus infection of the salivary glands of Frankliniella occidentalis, the western flower thrips"
