## Supplementary Figure S4 for "Sex-biased proteomic response to tomato spotted wilt virus infection of the salivary glands of *Frankliniella occidentalis,* the western flower thrips"

**Supplementary Figure S4:** Multi-level gene ontology (GO) analysis of saliva proteins of *F. occidentalis* males and females exposed (V) and non-exposed (NV) to TSWV. Numbers next to the GO terms indicate the number of proteins within the GO term. BP: Biological Process; MF: Molecular Function; CC: Cellular Component

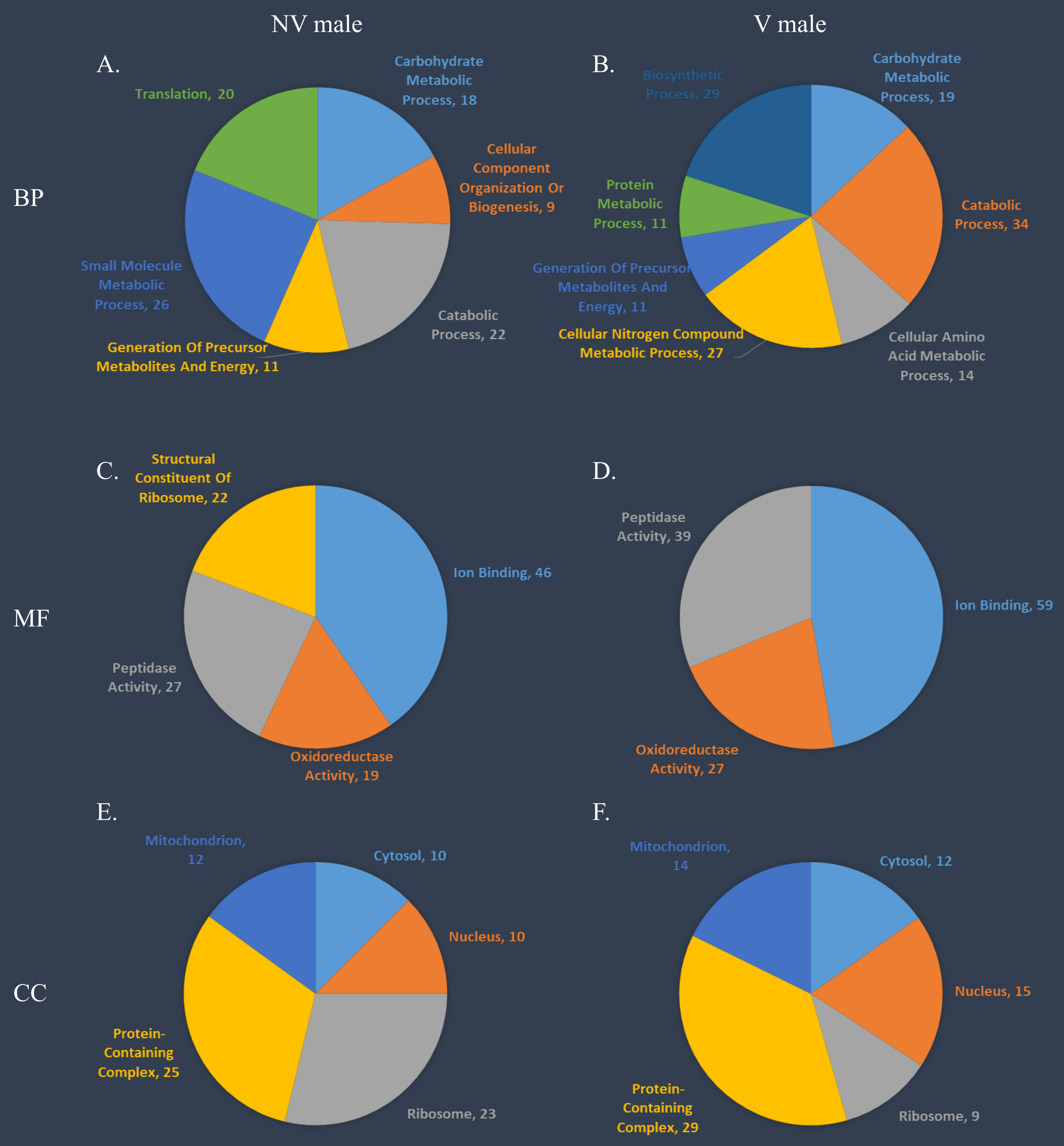

Rajarapu et al: "Sex-biased proteomic response to tomato spotted wilt virus infection of the salivary glands of *Frankliniella occidentalis*, the western flower thrips"

**Supplementary Figure S4, continued**

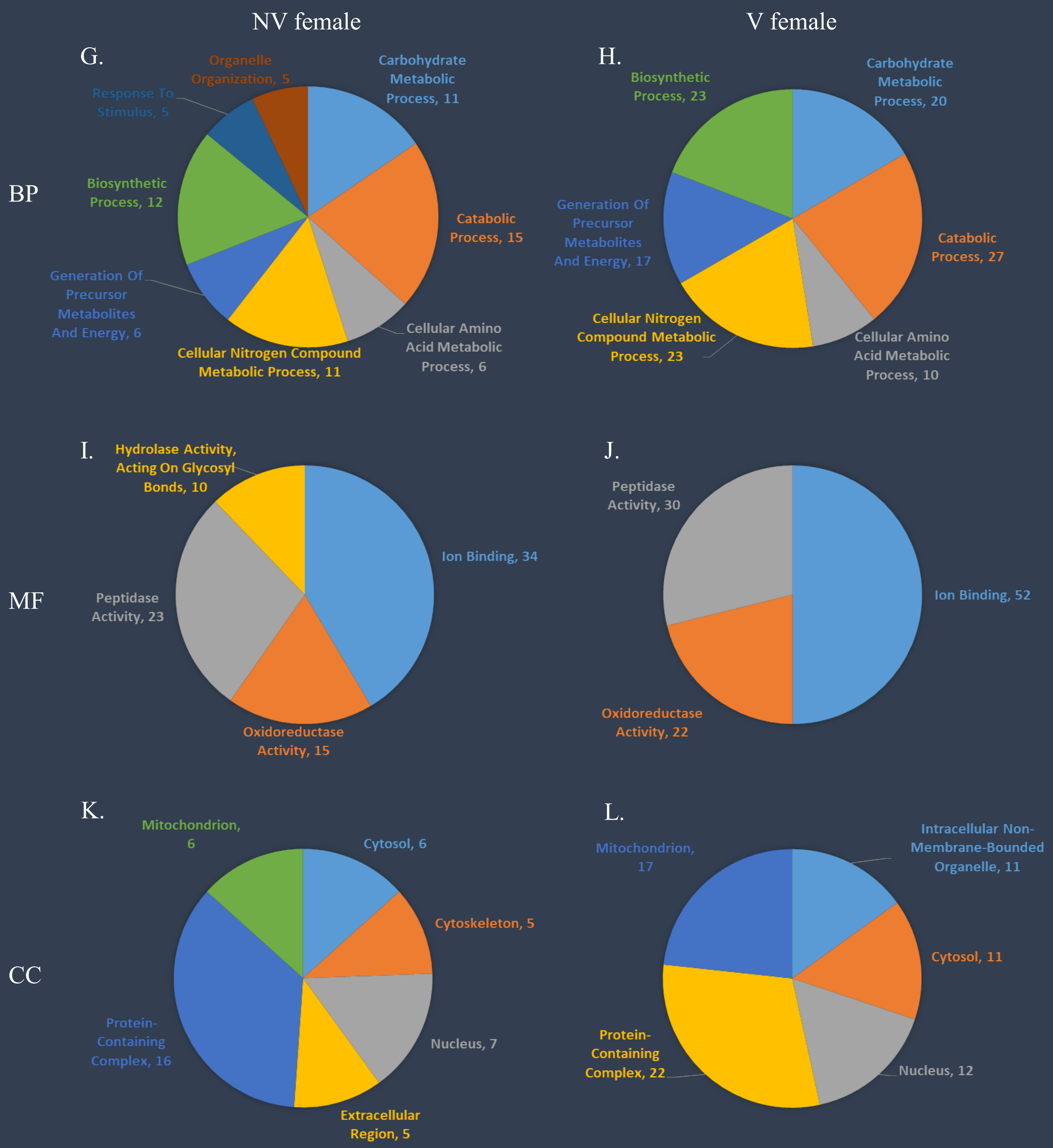
